# A Survey of the Kinome Pharmacopeia Reveals Multiple Scaffolds and Targets for the Development of Novel Anthelmintics

**DOI:** 10.1101/2020.08.20.259481

**Authors:** Jessica Knox, William Zuercher, Peter J. Roy

**Affiliations:** Department of Molecular Genetics, University of Toronto, Toronto, ON, M5S 1A8, Canada; The Donnelly Centre for Cellular and Biomolecular Research, University of Toronto, Toronto, ON, M5S 3E1, Canada; UNC Eshelman School of Pharmacy, University of North Carolina at Chapel Hill, NC, 27599, USA; Department of Pharmacology and Toxicology, University of Toronto, Toronto, ON, M5S 1A8, Canada

**Keywords:** C. elegans, EGFR, MEK1, PLK1, LET-23, MEK-2, PLK-1, EGF pathway, kinases, kinase inhibitors, ATP-binding pocket, nematodes, nematicides, anthelmintics

## Abstract

Over one billion people are currently infected with a parasitic nematode. Symptoms can include anemia, malnutrition, developmental delay, and in severe cases, death. Resistance is emerging to anthelmintic drugs used to treat nematode infection, prompting the need to develop new anthelmintics. Towards this end, we identified a set of kinases that may be targeted in a nematode-specific manner. We first screened 2040 inhibitors of vertebrate kinases for those that impair the model nematode *Caenorhabditis elegans*. By determining whether the terminal phenotype induced by each kinase inhibitor matched that of the predicted target mutant in *C. elegans*, we identified 17 druggable nematode kinase targets. Of these, we found that nematode EGFR, MEK1, and PLK1 kinases have diverged from vertebrates within their drug-binding pocket. For each of these targets, we identified small molecule scaffolds that may be further modified to develop nematode-specific inhibitors. Nematode EGFR, MEK1, and PLK1 therefore represent key targets for the development of new anthelmintic medicines.

**One sentence summary:** Druggable Kinases as Anthelmintic Targets

## Introduction

The burden of parasitic nematodes on humanity is severe. Over 1.5 billion people suffer from intestinal nematode infections alone [1, 2]. Chronic infection results in a myriad of symptoms including malnutrition, anemia and developmental delay. Left untreated, infections can lead to cognitive impairment that adversely affects education and employment and contributes to a cycle of poverty. Severe infections can lead to death [2, 3]. All told, parasitic nematode infections are responsible for an estimated disease burden of over 3.4 million disability-adjusted life years (DALYs), which is equivalent to that of tuberculosis or malaria [4, 5].

Anthelmintic drugs that are used to treat parasitic nematode infections include albendazole, mebendazole, diethylcarbamazine and ivermectin [3]. Emerging evidence indicates that mass drug administration is driving the evolution of resistance to these anthelmintics within human parasite populations [2]. Parasitic nematodes further impact our wellbeing by parasitizing livestock and crops, leading to billions of dollars in losses annually, increasing food costs and contributing to malnutrition [6-9]. As a result of intensive anthelmintic pressure, nematode parasites of livestock have rapidly developed resistance to every class of anthelmintic applied [10-14]. This reinforces the alarm raised by the emergence of resistance among the nematode parasites of humans. Hence, there is a clear need for the development of new compounds to combat parasitic nematodes of humans, animals, and plants.

Kinases hold potential for the development of novel anthelmintic drugs [15-22]. Kinases are a large family of enzymes that phosphorylate substrates to co-ordinate nearly every signaling pathway in the cell [23]. For example, upon binding of the epidermal growth factor (EGF) ligand, the EGF tyrosine kinase transmembrane receptor (EGFR) dimerizes and autophosphorylates itself, changing the conformation of its intracellular domain. Adaptor proteins are consequently recruited to initiate a RAS-driven MAP kinase signaling cascade that controls numerous events, including cell survival, proliferation, differentiation, growth, migration, and resistance to apoptosis [24, 25].

Here, we focus on kinases as potential anthelmintic targets for several reasons. First, there is extensive knowledge of kinase structure. The structures of protein kinases are among the most intensely studied of any protein family [26] (http://www.sgc.ox.ac.uk/research/kinases/); the Protein Data Bank (PDB) contains over 7000 kinase structures with and without inhibitors bound [27] (http://www.rcsb.org/pdb/). The vast number of solved kinase structures is likely due to their key roles in both development and pathogeneses, but also because of the relative ease of kinase purification and *in vitro* biochemical assays. Given a benchmark of 30% sequence identity for reliable prediction of a protein structure by homology modeling, models of many nematode kinases can be constructed based on the experimental 3D structure of their vertebrate orthologs [28, 29]. We reason that small differences between the respective orthologs’ drug-binding pockets may facilitate the development of nematode-specific kinase inhibitors.

Second, inhibitors against a vast array of kinases have been developed (reviewed in [30]). Indeed, vertebrate kinases are the second most targeted class of proteins for drug development after the G-protein coupled receptors [31]. Currently, there are 52 FDA approved small molecule inhibitors targeting protein kinases [32] and over 200 are undergoing clinical trials for a variety of indications including cancer, inflammation, and autoimmune diseases [33] (www.icoa.fr/pkidb/). Many more kinase inhibitors exist as pre-clinical research tools [30]. These include both ATP-competitive inhibitors and allosteric modulators. This well-established ‘druggability’ makes kinases attractive targets for anthelmintic exploration.

Finally, kinases are generally well-conserved. For example, the genome of the nematode *Caenorhabditis elegans* encodes 438 protein kinases and has close homologs of more than 80% of human kinases [34]. There is also a high degree of kinase conservation within the nematode phylum. For example, *C. elegans* has homologs of 95% of the kinases identified in the nematode parasite *Haemonchus contortus* [35]. Hence, drugs that target a particular vertebrate kinase may have utility against the respective ortholog in nematodes, and this kinase may be found broadly across the phylum.

Despite the conservation of kinases, there are sequence differences that distinguish nematode orthologs from their vertebrate counterparts. For example, the pairwise sequence similarity between homologous kinases of *H. contortus* and its sheep host *Ovis aries* is considerably lower (25% identity) than between *H. contortus* and *C. elegans* (35% identity) [35]. A similar trend is found when comparing sequence similarity of orthologous kinases from the human nematode parasite *Brugia malayi* to that of *C. elegans* versus that of its human host system [22]. These structural differences may be exploited to modify existing inhibitors of vertebrate kinases to derive nematode-specific inhibitors.

Here, we have identified candidate anthelmintic targets by exploiting the pharmacopeia of vertebrate kinase inhibitors. Our strategy consisted of three steps. First, we screened existing vertebrate kinase inhibitors for the ability to induce robust phenotypes in *C. elegans*. We focused on those inhibitors that yield phenotype that mimics the loss-of-function of the respective orthologous kinase in the worm. *C. elegans* kinases that satisfy this criterium are considered, at least preliminarily, as ‘druggable’. Second, for these druggable *C. elegans* kinases, we inspected whether the inhibitor binding site diverges from vertebrates in relevant residues using homology modelling. Third, for those *C. elegans* kinases that satisfy the first two criteria, we asked whether *C. elegans* residues that diverge from mammals are conserved within the nematode phylum. With this subset of kinases, it may be possible to modify the structure of the respective vertebrate kinase inhibitors to confer nematode-specificity. Finally, we used chemical genetic techniques to further investigate three of these druggable kinases that may be exploited to develop novel anthelmintics.

## Results

### A Screen of Vertebrate Kinase Inhibitors Reveals 17 Druggable Nematode Kinases

Towards identifying kinases that may serve as anthelmintic targets, we first screened 2040 vertebrate kinase inhibitors across four chemical libraries to identify those that disrupt the lifecycle of *C. elegans* (Figure 1; Supplementary File S1). 1160 of these molecules are kinase inhibitors that are used as either research tools or medicines and come from the DiscoveryProbe Kinase Inhibitor Library (APExBIO), the Ontario Institute for Cancer Research (OICR) Kinase Inhibitor Library and a subset of the Library of Pharmacologically Active Compounds (LOPAC, Sigma-Aldrich). We refer to these molecules as ‘commercial’ inhibitors. The remaining 880 molecules are from GlaxoSmithKline’s (GSK) Published Kinase Inhibitor Sets 1 and 2 (PKIS1 and PKIS2) [36-38]. GSK publicly released the PKIS library to foster basic research and academic drug discovery [37]. The core scaffolds in the PKIS were chosen based on previously published campaigns that targeted specific protein kinases, and these molecules were further screened against a panel of over 200 human kinases to assess specificity [36-38]. Hence, a preliminary assessment of the *in vitro* inhibitory profiles of the PKIS molecules is known and span a range of kinase selectivity profiles from narrow to broad spectrum. Many of the PKIS scaffolds are represented by multiple analogs within the library. Consequently, hits from the PKIS library can provide insight on structure-activity relationships (SAR). The four libraries that we screened contained a total of 1716 unique inhibitor structures.

**Figure 1.**
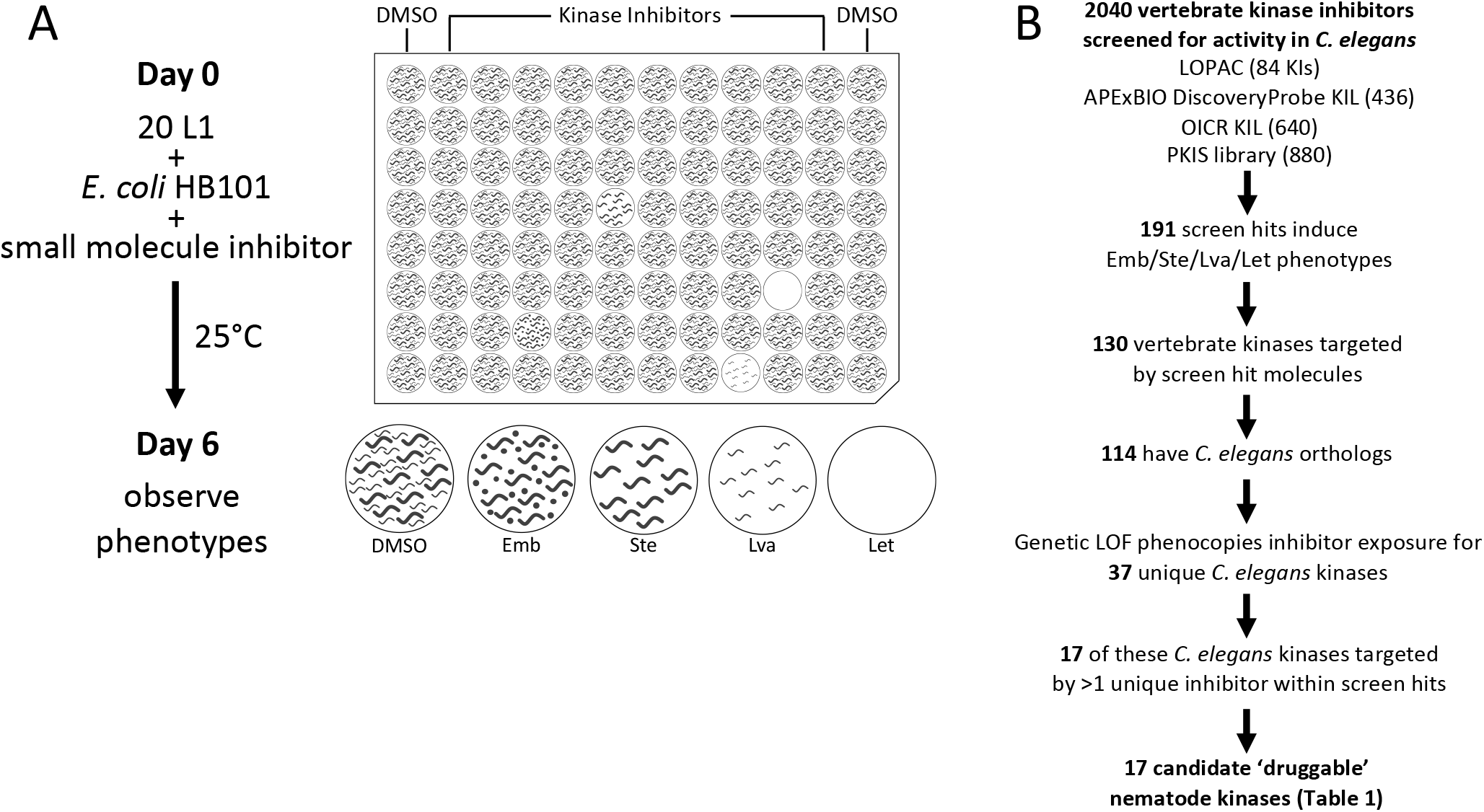
A screen of vertebrate kinase inhibitors reveals 17 druggable nematode kinases. (A) Schematic of small molecule screening methodology. (B) The pipeline used to identify candidate nematode kinase targets for structure analysis.

To identify druggable nematode kinases, we screened these kinase inhibitors for their ability to disrupt the viability, development and/or fecundity of *C. elegans*. We examined growth of the *C. elegans* culture after six days of co-incubation with the small molecules. We visually inspected the cultures to identify those compounds that induced any obvious phenotypes, including lethality (Let), larval arrest (Lva), sterility (Ste) and/or embryonic lethality (Emb) (Figure 1A). We screened each compound at a final concentration of 10 µM (OICR) or 60 µM (APExBIO, LOPAC, PKIS), which were technically convenient concentrations at which most molecules remain in solution. These concentrations may be considered high for *in vitro* and cell-based screening platforms, but are suitable for whole-animal *C. elegans* screens due to the nematode’s robust xenobiotic defenses [39, 40]. Using the criteria defined in the methods section, we identified 191 hits comprising 170 distinct inhibitors that reportedly target 130 vertebrate kinases that either kill, sterilize, or arrest the growth of *C. elegans* (Figure 1B; Supplementary File S1A; Supplementary Table 1).

**Table 1.** Druggable essential nematode kinase targets.

| Vertebrate Target | *C. elegans* Ortholog | # Hits | Exemplar Inhibitor<sup>a</sup> | Relevant Phenotypes<sup>b</sup> |
| --- | --- | --- | --- | --- |
| EGFR | LET-23 | 18 | Tyrphostin 23*<br>GSK306886A<br>GW576609B | Let Ste Rbs |
| ERBB2 |  | 6 |  |  |
| ERBB3 |  | 1 |  |  |
| ERBB4 |  | 9 |  |  |
| MAP2K1 | MEK-2 | 13 | Trametinib* | Emb Lva Let Ste Rbs |
| MAP2K2 |  | 13 |  |  |
| PLK1 | PLK-1<br>PLK-2 | 9 | MLN0905* | Emb Let Ste Rbs<br>Emb Ste Rbs |
| BRAF | LIN-45 | 14 | Dabrafenib (GSK2118436)* | Emb Let Ste |
| RAF1 |  | 12 |  |  |
| ARAF |  | 1 |  |  |
| AURKB | AIR-2 | 10 | SNS-314 Mesylate*<br>AMG-900* | Emb Let Ste Rbs |
| AURKA |  | 6 |  |  |
| AURKC |  | 4 |  |  |
| MTOR | LET-363 | 5 | Torin 1* | Emb Lva Let Ste |
| CDK1 | CDK-1 | 4 | Cdk1/2 Inhibitor III* | Emb Ste |
| MAPK11 | PMK-1 | 4 | SB-245392 | Let or Ste |
| MAPK14 |  | 3 |  |  |
| SRC | SRC-1 | 4 | PP2* | Emb Let |
| LCK |  | 2 |  |  |
| FYN |  | 1 |  |  |
| IGF1R | DAF-2 | 3 | GSK1838705A* | Let Ste Rbs |
| INSR |  | 2 |  |  |
| INSRR |  | 1 |  |  |
| STK10 | GCK-4 | 3 |  | Lva Let Rbs |
| CSNK1E | KIN-20 | 3 |  | Emb Let Ste Rbs |
| CSNK1D |  | 2 |  |  |
| GSK3A | GSK-3 | 2 | AZD2858* | Emb Let Ste |
| GSK3B |  | 2 |  |  |
| BUB1 | BUB-1 | 2 | GSK350559A | Emb Lva Let Ste |
| CSNK2A1 | KIN-3 | 2 | GW857175X | Emb Lva Ste |
| CSNK2A2 |  | 1 |  |  |
| CSNK1A1 | KIN-19 | 2 |  | Emb Ste |

<sup>a</sup> Commercially available inhibitors indicated with an asterisk (\*)
<sup>b</sup> Embryonic Lethal (Emb); Larval Arrest (Lva); Lethal (Let); Sterile (Ste); Reduced Brood Size (Rbs)

We next investigated whether these 191 vertebrate kinase inhibitors likely inhibit the orthologous *C. elegans* target or yield phenotype because of off-target effects. To do so, we asked whether the terminal phenotypes induced by a given kinase inhibitor phenocopied the genetic loss-of-function or RNAi phenotypes reported for the orthologous worm target on WormBase (WormBase web site, http://www.wormbase.org, release WS275, date 04-03-2020) [41]. We found that inhibitors induce phenotypes consistent with the genetic loss-of-function for 37 unique *C. elegans* kinases. We recognize that off-target phenotypes may be coincident with the anticipated phenotype and may confound interpretation at this point in the pipeline. However, downstream analyses (see below) further probe the relationship between the bioactive inhibitors and the anticipated targets.

To generate a higher-confidence list of druggable *C. elegans* kinases, we filtered the list of 37 kinases for those whose vertebrate orthologs were targeted by multiple structurally-distinct inhibitors that yielded the expected *C. elegans* loss-of-function phenotypes. In this way, we identified 17 higher-confidence druggable essential nematode kinase targets (Figure 1B; Table 1). Regardless of the ultimate anthelmintic utility of these 17 druggable kinases, the respective inhibitors may be useful chemical tools for the community, allowing for the temporal control of kinase activity and simplifying the investigation of these essential cellular components in *C. elegans*.

### Three Kinases are Candidate Targets for Anthelmintic Development

We investigated whether any of the 17 higher-confidence kinase targets exhibit sequence divergence between nematodes and vertebrates within the drug-binding pocket. This divergence could then be exploited to develop nematode-specific inhibitors. To do this, we first used the SWISS-MODEL pipeline to generate protein homology models, mapping the *C. elegans* primary sequence onto the experimentally derived 3D crystal structure of the respective vertebrate kinase ortholog [42]. These homology models were visualized using PyMOL (The PyMOL Molecular Graphics System, Version 2.1.1, Schrödinger, LLC).

To test the validity of a homology modeling approach, we first created a homology model of the *C. elegans* LET-23 kinase domain based on the solved structure of the orthologous human EGFR kinase domain (Protein Data Bank (PDB) structure 2ITX; [43]) (Supplementary Figure 1A). We then compared the LET-23 homology model to the experimentally derived crystal structure of the LET-23 kinase domain (PDB: 5WNO [44]; Supplementary Figure 1B). We found that the side chains of the residues that line the ATP-binding (and drug-binding) pocket are similarly oriented towards the active site in both models. This analysis supports the utility of the homology modeling approach to compare presumptive drug binding pockets of nematode kinases to orthologous vertebrate structures.

For each of the 17 druggable nematode kinases, we examined whether any residues that line the drug-binding pocket of the *C. elegans* kinase diverge from their respective vertebrate ortholog. Particular attention was paid to those residues that reside within 5Å of the bound inhibitor (Supplementary Figure 2). To determine whether the pocket-lining residues that diverge in *C. elegans* also diverge within diverse parasitic nematode species, we generated and analyzed multiple sequence alignments. This pipeline revealed three potential anthelmintic targets that include EGFR (LET-23 in *C. elegans*), the MAP kinase kinase MEK1 (MEK-2 in *C. elegans*), and polo-like kinase PLK1 (PLK-1 in *C. elegans*) (Figure 2). Structural modeling of these three kinase domains from the human parasitic nematode *B. malayi* confirmed that the divergent residues line the prospective drug binding pockets and provide an opportunity to develop nematode-specific kinase inhibitors (Supplementary Figure 2).

**Figure 2.**
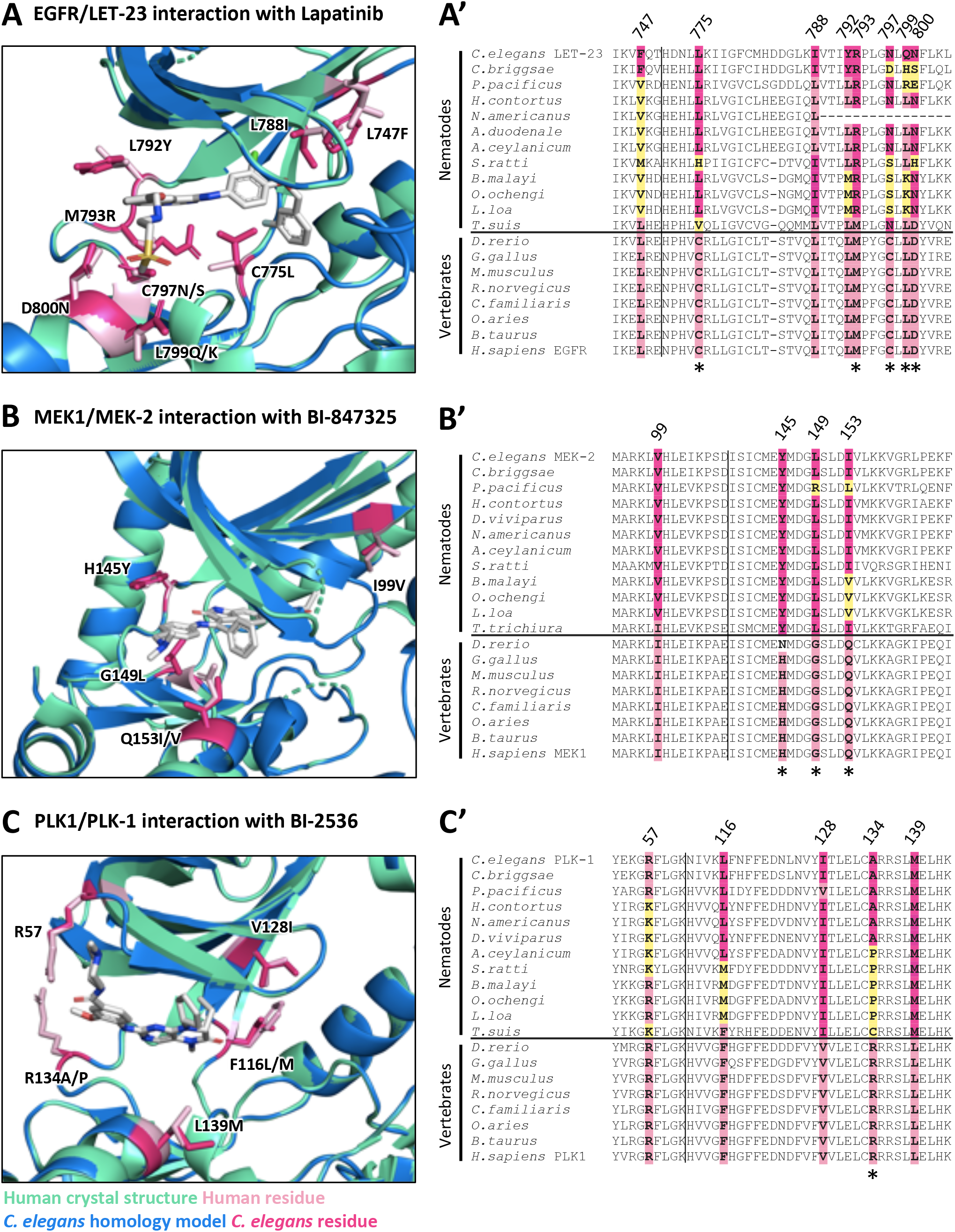
Structure analysis identifies LET-23, MEK-2 and PLK-1 as candidate targets for anthelmintic development. *C. elegans* homology models (blue) for LET-23, MEK-2 and PLK-1 aligned with human crystal structure (green) for EGFR (PDB: 1XKK, [81]), MEK1 (PDB: 5EYM, [60]) and PLK1 (PDB: 2RKU, [65]) are shown in A, B and C respectively. Key divergent residues proximal to the inhibitor binding sites are highlighted in dark pink (*C. elegans* residue) and light pink (human residue) within the structure diagrams (A,B,C). Residues are labeled according to their position in the human kinase, with the first letter indicating the identity of the vertebrate residue and the latter indicating the identity of the nematode residue(s). The conservation of these residues among free-living nematodes, parasitic nematodes and vertebrates is displayed in the corresponding sequence alignments (A’, B’, C’). Yellow residues in the sequence alignments highlight those nematode residues that differ in identity from both vertebrate and *C. elegans* sequence at the location of these divergent residues of interest. Residues that have distinct physicochemical properties between vertebrates and nematodes are indicated with an asterisk below the alignment.

### LET-23 Divergence May Interfere with its Ability to Interact with EGFR Inhibitors

Our pipeline revealed LET-23, the *C. elegans* EGFR ortholog, as a candidate anthelmintic target. LET-23 and its conserved signaling pathway (Figure 3) regulates a wide variety of developmental processes including vulval induction and excretory duct cell formation and is one of the best understood receptor tyrosine kinases in *C. elegans* [24, 25, 45, 46]. LET-23, which is expressed in the presumptive vulval epithelial cells, responds to LIN-3/EGF that is secreted by the nearby anchor cell to coordinate the development of the vulva tissue [47]. Too little signaling through LET-23 renders cells incapable of developing vulva tissue, resulting in what is known as a vulvaless (Vul) phenotype [48]. LET-23 also plays a Ras-independent role in promoting ovulation through the PLCγ-IP_3_ pathway, resulting in a sterile (Ste) phenotype upon loss of LET-23 function [25]. Weak *let-23* mutants such as *let-23(sy1)* are vulvaless but not sterile. Consequently, *let-23(sy1)* adults are filled with internally hatched embryos that destroy their hermaphrodite mothers, which is a phenotype known as bag-of-worms (Bag). When LET-23 pathway signalling is overactive, too many cells develop into primary vulval tissue, creating a multivulva (Muv) phenotype [49]. LIN-3/LET-23 signaling via Ras-ERK also plays an integral role in excretory duct cell fate specification, morphogenesis and differentiation [50, 51]. Proper development of the excretory duct cell is required for osmoregulation [52, 53]. Homozygous null *let-23* animals are therefore larval lethal and die with a distinctive rod-like, fluid-filled appearance due to the loss of the excretory duct cell [50].

**Figure 3.**
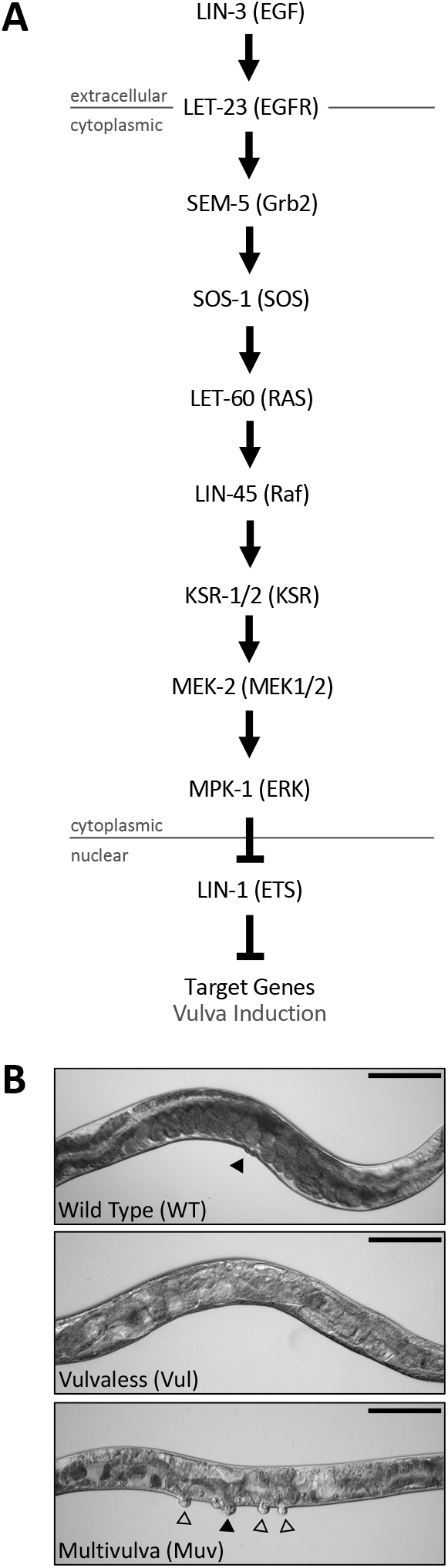
The conserved *C. elegans* EGFR/Ras/MAPK pathway controls vulval induction signaling. (A) A schematic of the conserved EGFR signaling pathway in nematodes. The vertebrate orthologs for each pathway component are shown in brackets. Changes in signaling levels through this pathway result in the Vulvaless (Vul) and Multivulva (Muv) vulva induction phenotypes shown in (B). Filled arrows indicate primary functional vulva, clear arrows indicate ectopic vulval protrusions. Scale bar 0.1 mm.

Our collection of kinase inhibitors include 66 distinct commercial inhibitors of EGFR or paralogous ERBBs (Supplementary File S2A). Of these, only two (Tyrphostin AG879 and Tyrphostin 23) induce robust phenotypes in *C. elegans*. One possibility for why so few of these inhibitors induce *let-23* phenotypes is that they may have unfavorable ADME properties in nematodes and fail to reach target. We tested this possibility by asking whether one of these molecules, gefitnib, could induce *let-23*-like phenotypes in a humanized strain of *C. elegans* in which the kinase domain of LET-23 is replaced by that from human EGFR. This strain was kind gift from Dr. Jaegal Shim [54]. Previous work demonstrated that the humanized LET-23 could rescue the vulvaless phenotype of the *let-23(sy1)* mutant, which we confirmed (Figure 4). Gefitnib failed to induce a vulvaless phenotype in wild type animals, but did induce a vulvaless phenotype in the humanized strain (Figure 4). This data indicates that an ADME barrier is not the reason why gefitnib, and perhaps other EGFR/ERBB inhibitors, fail to inhibit worm LET-23.

**Figure 4.**
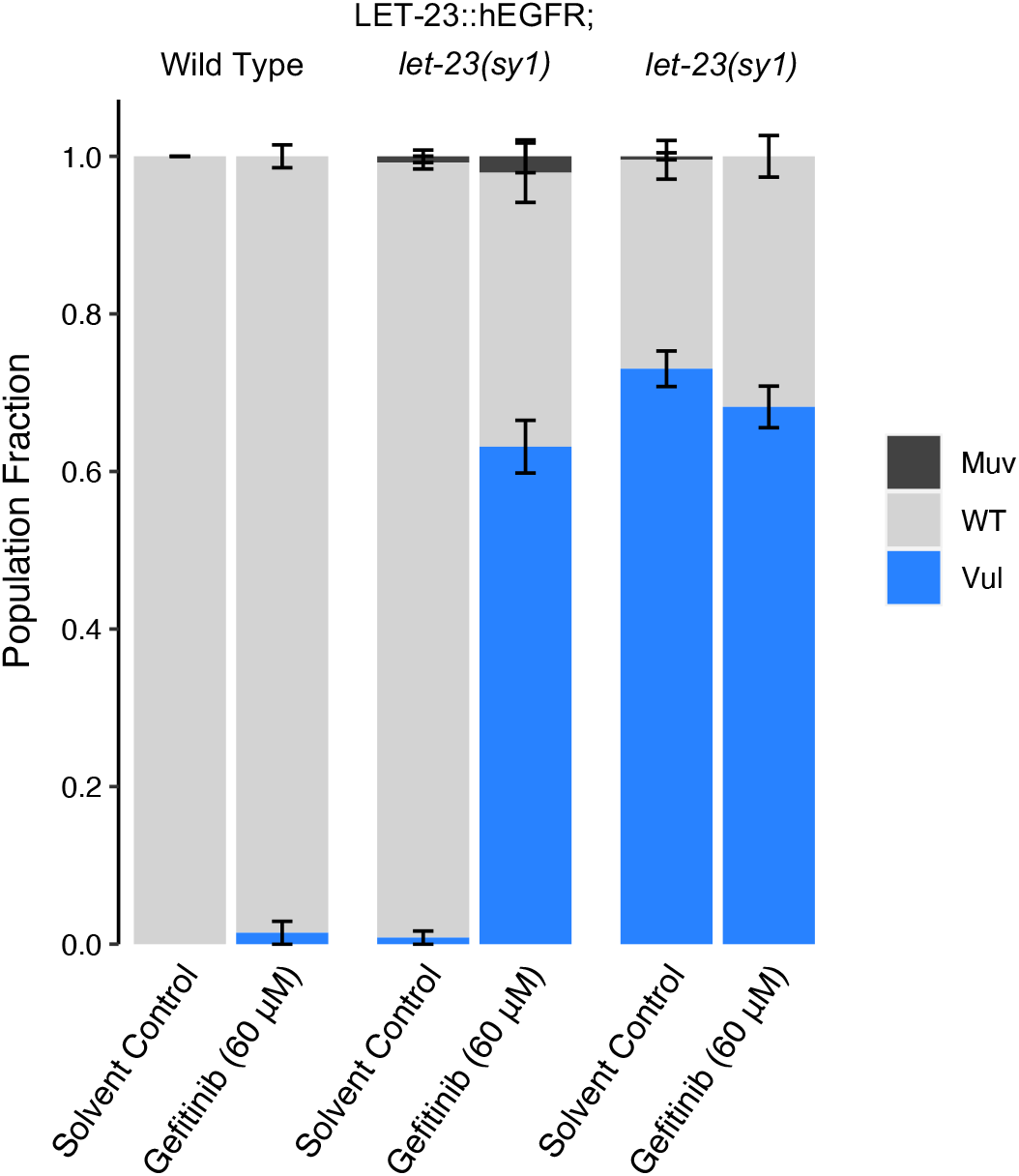
Structural divergence in EGFR/LET-23 impacts response to EGFR inhibitor Gefitinib. The Vulvaless phenotype observed in the *let-23(sy1)* mutant is rescued by the expression of a chimeric protein whereby the kinase domain of LET-23 is replaced with human EGFR (LET-23::hEGFR). Gefitnib induces a vulvaless phenotype in the humanized strain, but not in wild type animals. DMSO is used as the solvent control. Error bars indicate SEM.

A second possibility for why so few EGFR/ERBB inhibitors disrupt LET-23 is that there may be structural differences within the drug-binding pocket of LET-23 that hinders interaction with EGFR/ERBB inhibitors. We investigated this possibility by comparing the structure of EGFR’s drug binding site to that of *C. elegans* LET-23 (as described above-see Supplementary Figure 2A). Overall, there is 44% identity in sequence in the kinase domain between LET-23 and human EGFR. Within the drug-binding pockets of the human and *C. elegans* structures, we found 10 obvious amino acid differences (Supplementary Figure 2A), eight of which also vary between vertebrate hosts and parasitic species (Figure 2A’). Five of these eight residues have distinct physicochemical properties from the vertebrate residues and could account for differential compound binding (Figure 2A’). We conclude that the structural differences within the presumptive drug-binding pocket of LET-23 likely accounts for the inability of the EGFR/ERBB inhibitors to induce *let-23* phenotypes. It is these structural differences that may allow for the rationale design of nematode-specific EGFR inhibitors.

### Three Scaffolds for the Development of Nematode-Specific LET-23 Inhibitors

We identified 20 PKIS inhibitors that induce *let-23*-like hypomorphic phenotypes in *C. elegans* (Figure 5A) and show some degree of specificity for EGFR/ERBB inhibition (Supplementary File S2B) [36-38]. These compounds fall within three related core scaffolds: the <u>4</u>-<u>a</u>nilino <u>q</u>uinazolines (4AQs); a 4AQ derivative scaffold called the <u>q</u>uinazoline <u>b</u>enzimidazoles (QBIs); and the <u>a</u>nilino <u>t</u>hien<u>op</u>yrimidines (ATOPs) (Figure 5B). The quantities of these compounds did not allow for extensive follow up experiments. However, we were able to test one exemplar from each scaffold (GW576609B (a 4AQ), GW272142A (a QBI), and GSK306886A (an ATOP)) in dose-response analyses, which revealed additional *let-23*-like phenotypes that include Bag and sterility (Figure 5C). These phenotypes provide further confidence that the 4AQs, QBIs and ATOPs likely disrupt LET-23 function.

**Figure 5.**
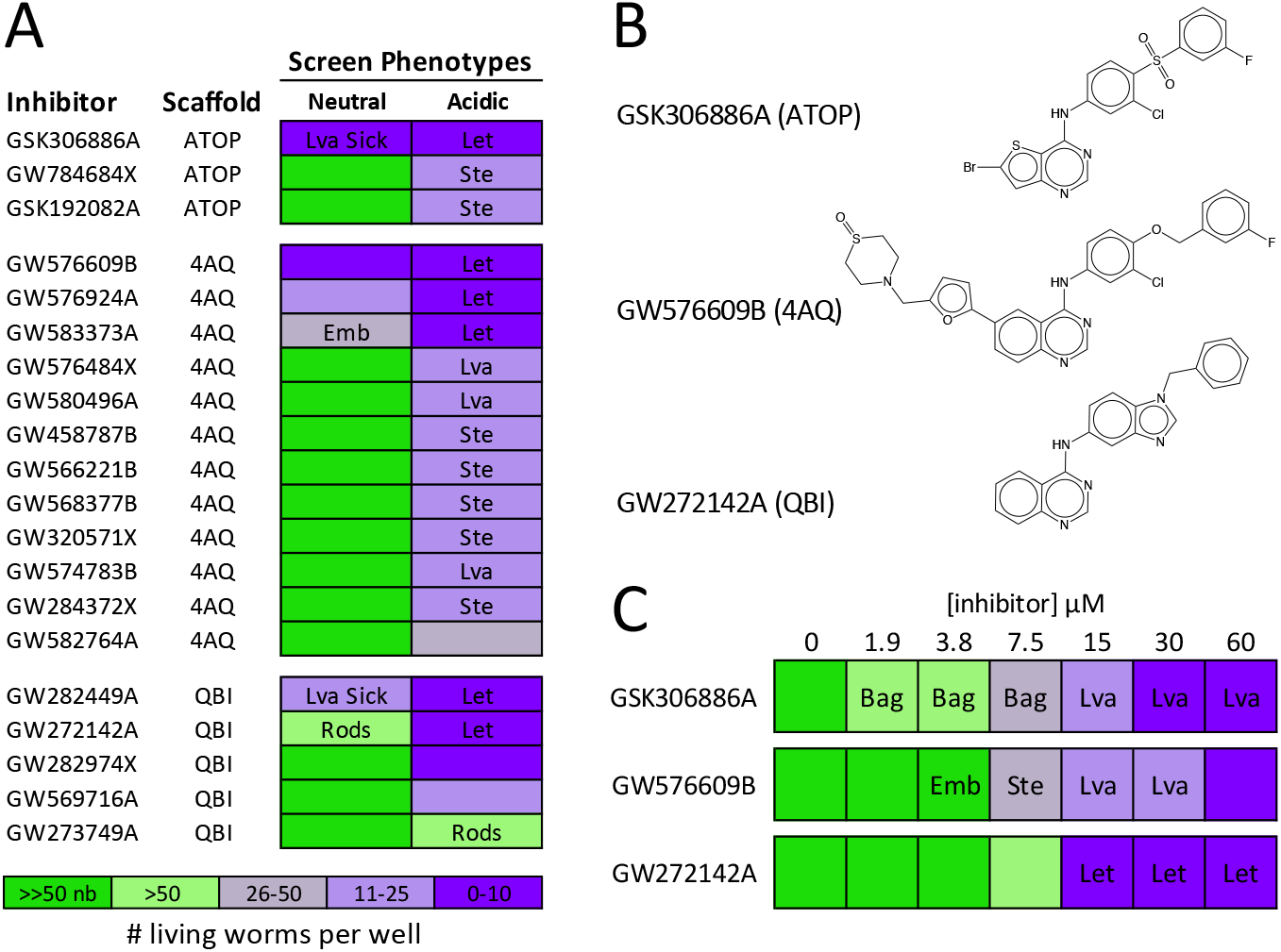
Three scaffolds induce LET-23 loss-of-function phenotypes in *C. elegans*. (A) The structurally related 4-anilino quinazolines (4AQ), anilino thienopyrimidines (ATOP) and quinazoline benzimidazoles (QBI) scaffolds induce LET-23 loss-of-function phenotypes in our *C. elegans* chemical screens. (B) An example of a worm-active structure from each scaffold is displayed. (C) Dose response analyses reveal phenotypes relevant to EGFR/MAPK pathway inhibition in *C. elegans* including sterility (Ste) and the bag-of-worms phenotype (Bag). Additional phenotypes including embryonic lethality (Emb) and larval arrest (Lva) are shown and the resulting population growth defects are indicated by the colour coded scale (nb, no bacteria remaining in the well). Dose response analyses for GSK306886A and GW576609B were performed in neutral media, GW272142A analysis was performed in acidic media.

By analysing the bioactivity of 4AQ-based structures in *C. elegans*, we were able to complete a small SAR analysis (Supplementary Figure 3). We found several substructural features that may improve activity in *C. elegans*, including a thiomorpholine 1-oxide ring structure in the R_1_ position, a furan as opposed to a thiazole as the X_1-3_ substituted ring and a halogen in the R_5_ position. These features are embodied in GW576609B, which could serve as a key scaffold on which to base the rational design of a nematode-specific EGFR inhibitor. An insufficient number of QBIs and ATOPs precluded us from performing a similar SAR analysis on these scaffolds.

### MEK-2 is a Druggable Kinase with Conserved Divergence Among Nematodes

Our pipeline revealed worm MEK-2 as a candidate anthelmintic target. MEK-2 is a MAP kinase kinase ortholog of mammalian MEK1/2 that functions downstream of the EGF receptor LET-23 in *C. elegans* (Figure 3) [25, 55, 56]. The spectrum of worm *mek-2* loss-of-function phenotypes is similar to that *of let-23* mutants; *mek-2* loss-of-function mutants die as rod-like sticks while weaker reduction-of-function mutants that escape lethality become sterile and/or vulvaless [55].

The kinase domain of worm MEK-2 is 60% identical to vertebrate MEK1. Much of the ATP/drug binding pocket of nematode MEK-2 is identical to vertebrate MEK1. However, there are three divergent residues with distinct properties on the hinge that connect the N and C lobes (Figure 2B, Supplementary Figure 2B). The divergence is well-conserved within the nematode phylum (Figure 2B’). Compounds that interact with MEK1/2 near the hinge may therefore have the potential to be modified for increased nematode specificity over the vertebrate ortholog.

In our initial survey, we screened 18 unique commercial inhibitors of vertebrate MEK1/2 (Figure 6). Nine of these inhibitors dramatically slowed population growth and/or induced *mek-2*-like hypomorphic phenotypes, including bag-of-worms, sterility, and embryonic lethality (Figure 6A-A’’). This activity suggested that MEK-2 is a druggable target.

**Figure 6.**
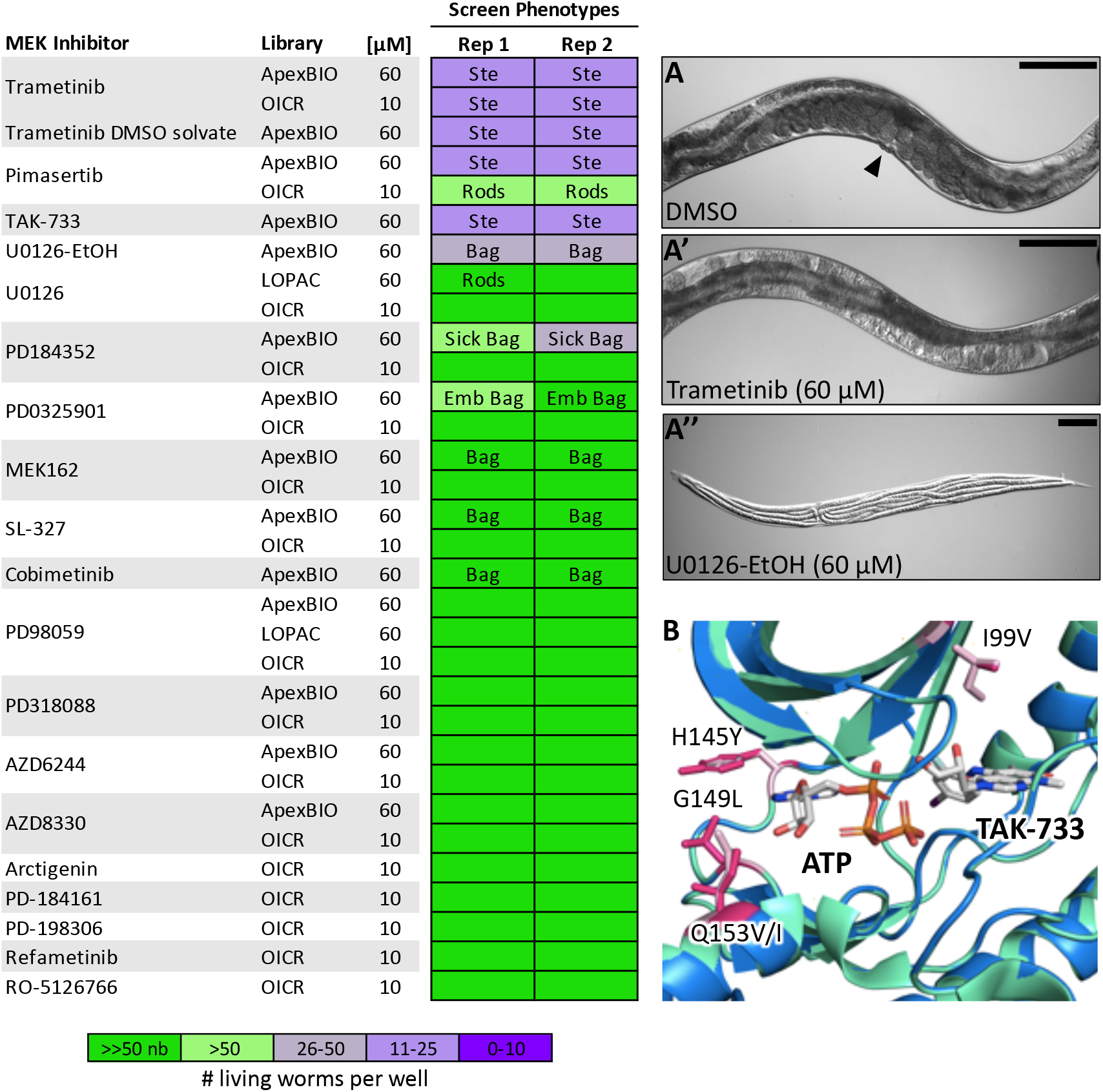
Allosteric MEK inhibitors induce sterility, embryonic lethality and vulva development phenotypes in *C. elegans*. 18 unique MEK inhibitors from the commercial libraries screened were included in our screen. These inhibitors were enriched for hits that induce the expected MEK-2 loss-of-function phenotypes including sterility (Ste) and the bag-of-worms phenotype (Bag), resulting from vulval induction defects preventing egg-laying (A-A’, scale bar 0.1 mm). The resulting population growth defects are indicated by the colour coded scale (nb, no bacteria remaining in the well). (B) The allosteric inhibitor binding site of *C. elegans* MEK-2 (in blue) is well conserved with that of vertebrate MEK1 (in green, seen co-crystalized with allosteric inhibitor TAK-733 (PBD: 3PPI). The allosteric site contains only one amino acid difference, highlighted in pink (I99V).

To validate worm MEK-2 as the target of these vertebrate MEK1/2 inhibitors *in vivo* we took a chemical-genetic approach using three structurally distinct worm-active compounds, trametinib, pimasertib, and U0126-EtOH. LET-60/RAS functions upstream of MEK-2 (Figure 3) [55, 56]. A gain-of-function allele of *let-60(n1046)* induces a partially penetrant multivulva (Muv) phenotype [57]. If a compound inhibits MEK-2, it should suppress the Muv phenotype of *let-60(n1046)*. By contrast, the LIN-1 ETS transcription factor is a negative regulator of the EGFR/MAPK pathway. A loss-of-function allele of *lin-1(e1275)* also results in a Muv phenotype. However, MEK-2 functions upstream of LIN-1 [58]. Hence, molecules that disrupt MEK-2 should not suppress the MUV phenotype of *lin-1(e1275)* animals. We found that all three MEK1/2 inhibitors suppressed the Muv phenotype of *let-60(n1046)* mutants at sub-micromolar concentrations and failed to suppress the Muv phenotype of *lin-1(e1275)* mutants up to the highest concentrations tested (120 µM) (Figure 7A-C). These data indicate that the MEK1/2 inhibitors likely act on their expected target *in vivo* and confirms that MEK-2 is a druggable target in nematodes.

**Figure 7.**
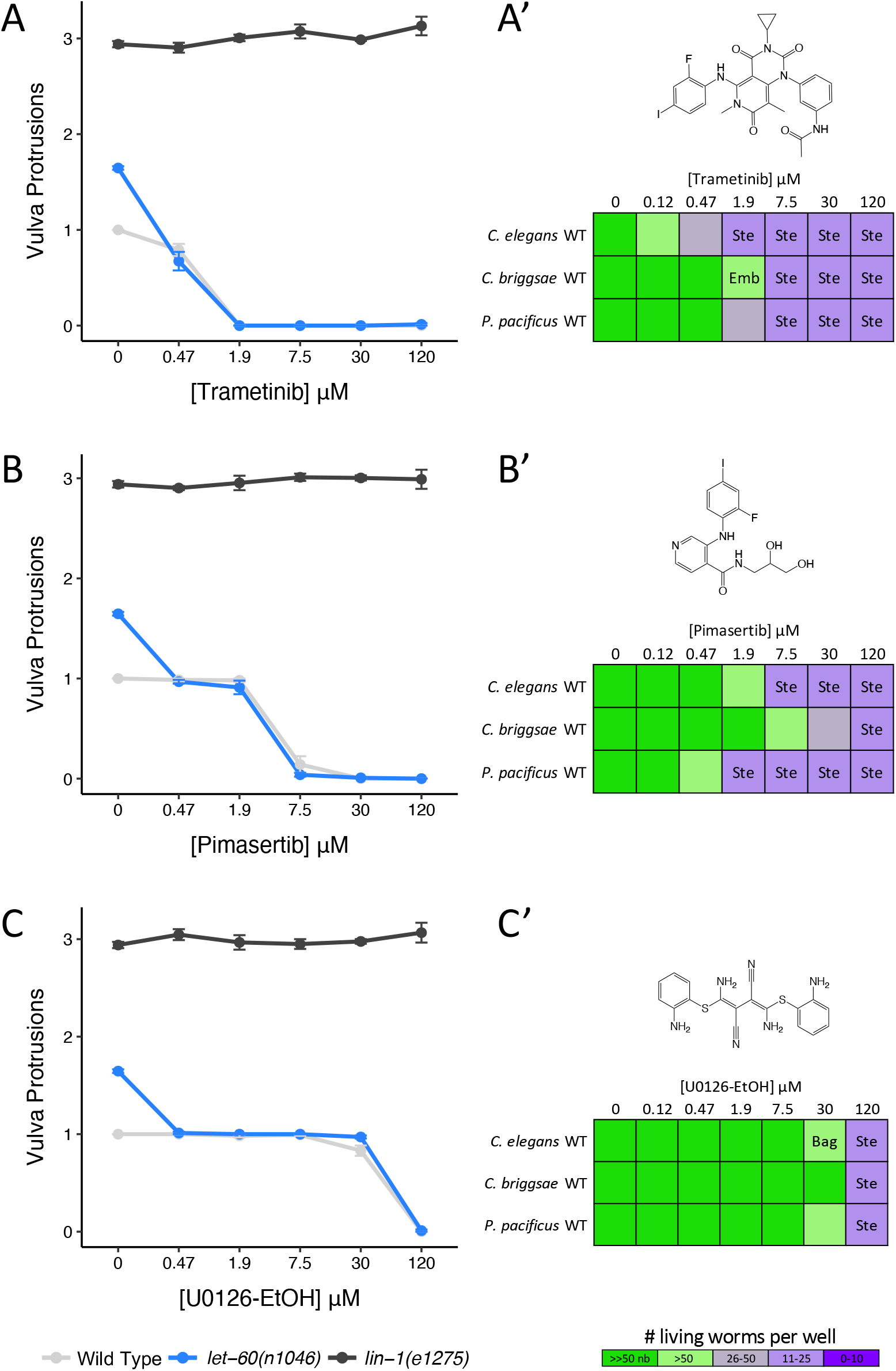
Vertebrate allosteric MEK inhibitors target worm MEK-2, inducing loss-of-function phenotypes across nematode species. (A-C) Three structurally distinct allosteric MEK inhibitors suppress the Multivulva phenotype of upstream *let-60(n1046)* gain-of-function mutants, but have no effect on the Multivulva phenotype in downstream *lin-1(e1275)* loss-of-function mutants. The average number of vulva protrusions observed per worm in each condition quantified over 3 biological replicates is shown. Error bars indicate SEM. (A’-C’) Phenotypes induced by allosteric MEK inhibitors in free-living nematode species *C. elegans, C. briggsae* and *P. pacificus*. Sterility (Ste), embryonic lethality (Emb) and bag-of-worms (Bag) phenotypes are reported along with the resulting population growth defects indicated by the colour coded scale (nb, no bacteria remaining in the well).

Finally, we examined whether the three aforementioned allosteric MEK inhibitors induce *mek-2*-related loss-of-function phenotypes in two additional free-living nematode species, *C. briggsae* and *Pristionchus pacificus*. We found that these inhibitors induce the expected phenotypes across all three nematode species in a dose-dependent manner with trametinib being the most potent (Figure 7A’-C’). Together, these results show that MEK-2 may be an excellent anthelmintic target.

### Divergence of the MEK-2 Hinge May Be Exploited to Develop Nematode-Specific Inhibitors

The commercial MEK1/2 inhibitors that we screened are allosteric modulators that do not compete for the ATP binding pocket (Figure 6). Instead, they bind near the base of the αC helix, which is distal to the divergent nematode sequence in the hinge region (Figure 6B; PDB: 3PP1 [59]). It is therefore unlikely that these allosteric MEK inhibitors can be modified to be nematode-specific. By contrast, MEK1/2 ATP-competitive inhibitors may be far better scaffolds upon which to build nematode-specific analogs because the hinge that lines the ATP-binding pocket of nematode MEK-2 is divergent from that of vertebrate MEK1/2 (Figure 6B).

We investigated whether the differences in nematode MEK1/2 structure are likely sufficient to confer phylum-specificity. We first inspected the co-crystal structure of an ATP-competitive inhibitor of MEK1/2 called BI-847325 (PDB: 5EYM) [60], which binds to the vertebrate MEK1/2 close to the hinge region (Figure 2B). Inspection of the MEK-2 homology model reveals that the leucine of the nematode sequence (G149L) may sterically hinder BI-847325’s terminal amine and prevent binding (Figure 2B). Of note, BI-847325 is one of the only established MEK1/2 specific inhibitors that competes for the ATP-binding pocket; the majority of MEK1/2 inhibitors are non-ATP competitive allosteric modulators [61].

We tested the hypothesis that BI-847325 is unable to disrupt MEK-2 activity *in vivo* through a dose-response analysis. BI-847325 inhibits vulval induction at 120 µM in wild type animals (Figure 8A). If BI-847325 reduced vulval induction through the inhibition of MEK-2, then BI-847325 should suppress the multivulva (Muv) phenotype of the upstream RAS gain-of-function mutation (*let-60(n1046)*), but not the Muv phenotype of the down-stream ETS transcription factor loss-of-function mutation (*lin-1(e1275)*). Instead, we found that BI-847325 reduced vulval induction in both mutants, suggesting that BI-847325 can access target in the worm, but is modulating a target other than MEK-2 (Figure 8A). BI-847325 is a dual inhibitor of vertebrate MEK and Aurora kinases [60]. Reduced vulva induction by BI-847325 may be due to the inhibition of a *C. elegans* Aurora kinase. Indeed, loss of function of AIR-2 Aurora kinase ortholog in the worm results in vulvaless, embryonic lethality and sterility phenotypes in *C. elegans* [62], which are all phenotypes induced by BI-847325 (Figure 8A’). Furthermore, there is no sequence divergence in the drug binding pocket of worm AIR-2 (relative to human Aurora kinases) that would impede inhibitor binding (Supplementary Figure 2E) [60]. We conclude that BI-847325 is bioavailable to the worm but is unlikely to engage MEK-2. Hence, the divergence in the hinge region of worm MEK-2 may allow for the rational design of a nematode-specific ATP-competitive MEK inhibitor. The core scaffold of BI-847325 would be an appropriate starting point for such an effort.

**Figure 8.**
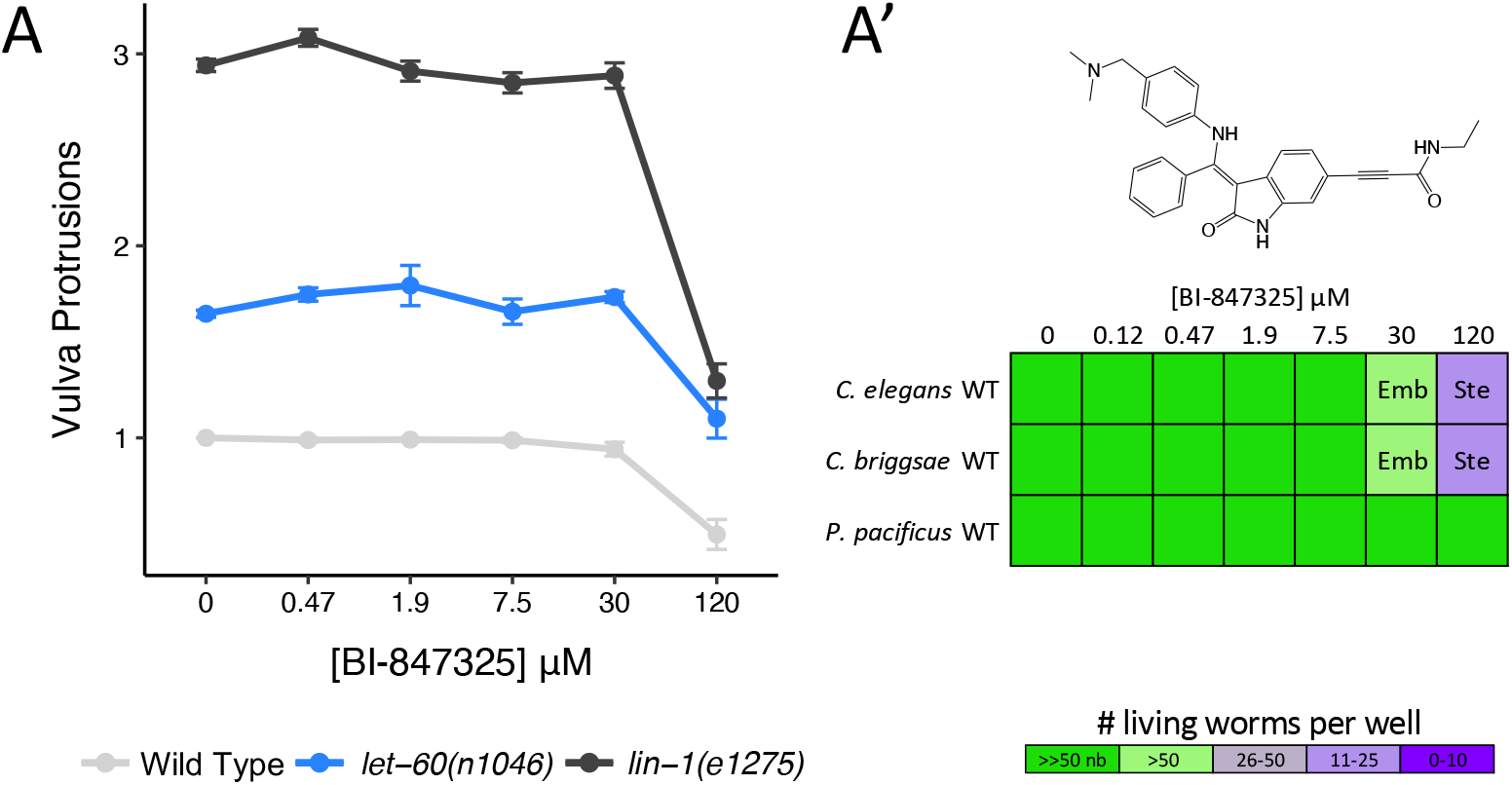
ATP-competitive MEK inhibitor BI-847325 does not engage nematode MEK-2. (A) The ATP-competitive MEK inhibitor BI-847325 significantly suppresses vulva induction in Wild Type and both mutants at the highest concentration tested (120 μM) relative to solvent controls (Student’s T-test: p<0.001). The average number of vulva protrusions observed per worm in each condition quantified over 3 biological replicates is shown. Error bars indicate SEM. (A’) BI-847325 induces sterility (Ste) and embryonic lethality (Emb) phenotypes in *C. elegans* and *C. briggsae* resulting in the population growth defects indicated by the colour coded scale (nb, no bacteria remaining in the well).

### PLK-1 is a High Priority Target for Anthelmintic Development

The *C. elegans* Polo-Like Kinase 1 ortholog PLK-1 regulates the meiotic cell cycle and embryonic polarity in the worm. *C. elegans* PLK-1 is 64% identical in the kinase domain to human PLK1. Loss-of-function mutations in *plk-1* result in embryonic lethality and sterility phenotypes, along with a protruding vulva phenotype [63, 64]. PLK1 inhibitors from three core scaffolds within the PKIS library induce phenotypes with a strong correlation to those observed upon genetic loss of *plk-1*. These include the benzimidazole N-thiophenes (BTs), the 2,4-dianilino pyrrolopyrimidines, and the 2,4-dianilino pyrimidines (Figure 9; Supplementary File S2D). The strong correlation between the established *plk-1* mutant phenotypes and those caused by the PLK1 inhibitors across the three scaffolds gave us confidence in the potential of PLK-1 as a druggable target in the worm.

**Figure 9.**
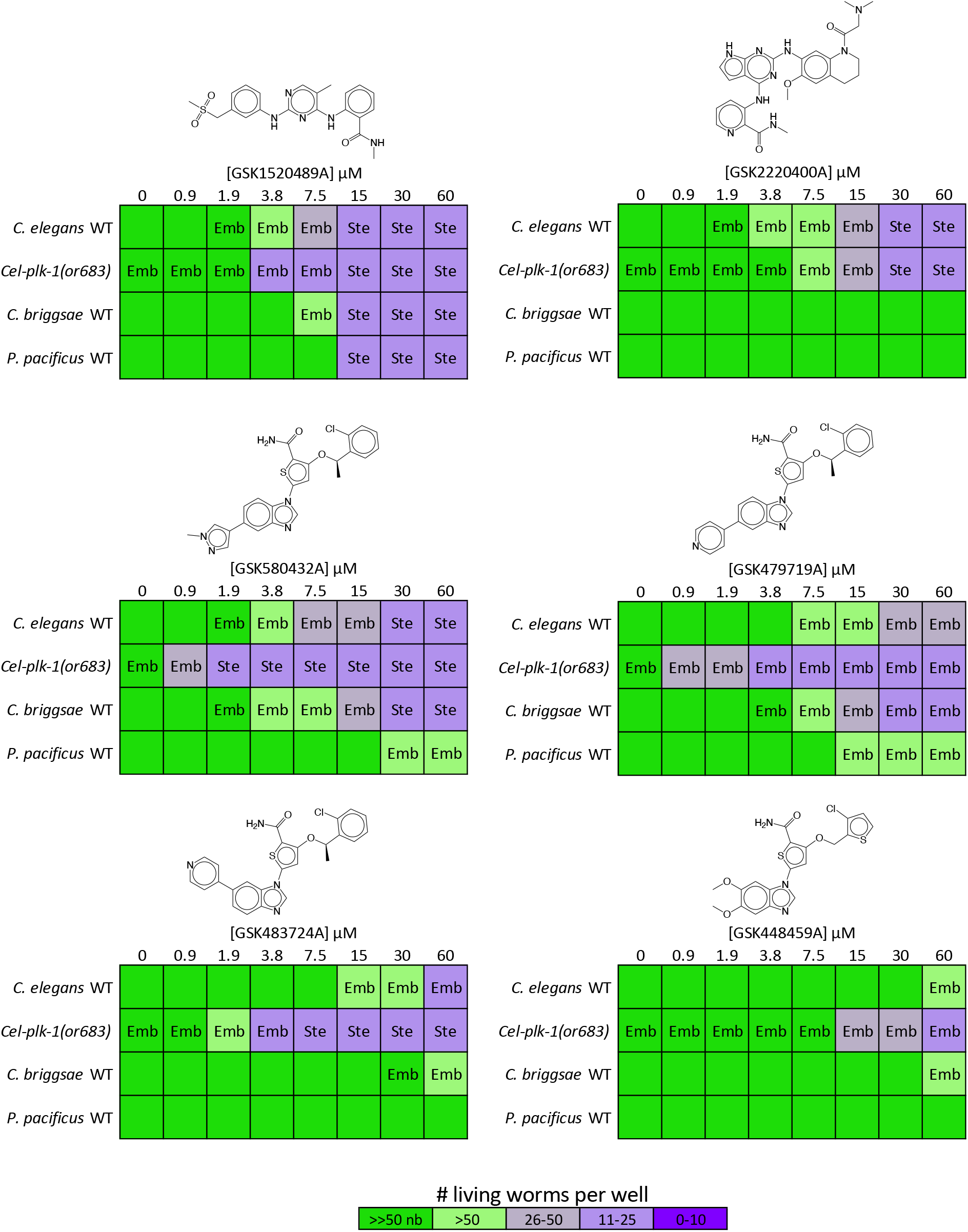
PLK1 inhibitors induce embryonic lethality and sterility phenotypes in nematodes. PLK1 inhibitors from three core scaffolds including the 2,4-Dianilinopyrimidines (GSK1520489A), the 2,4-Dianilino pyrrolopyrimidines (GSK2220400A) and Benzimidazole N-thiophenes (GSK580432A, GSK479719A, GSK483724A, and GSK448459A) induce phenotypes consistent with loss of *C. elegans* PLK-1 including sterility (Ste) and embryonic lethality (Emb) resulting in the population growth defects indicated by the colour coded scale (nb, no bacteria remaining in the well).

We modeled the protein structure of *C. elegans* PLK-1 against the crystal structure of human PLK1 (PDB:2RKU, [65]). Four residues in the ATP binding pocket differ between *C. elegans* PLK-1 and human PLK1 (Figure 2C, Supplementary Figure 2C). A multiple sequence alignment with additional nematode and vertebrate species shows that nematode residues differ at each of these locations, with a greater degree of conservation with the specific *C. elegans* amino acid among the parasitic Clade V nematodes (Figure 2C’). A fifth residue of interest was identified (R57) that is conserved between *C. elegans* and vertebrates but differs in a number of the parasitic nematode sequences examined (Figure 2C’).

We retested a selection of available PKIS compounds from the three core scaffolds of interest in *C. elegans* and two additional free-living nematode species, *C. briggsae* and *P. pacificus* (Figure 9). We found that the 2,4-dianilinopyrimidine GSK1520489 and the benzimidazole thiophene (BT) inhibitors GSK580432 and GSK479719 induced relevant embryonic lethality and/or sterility phenotypes across all species tested. To test whether these compounds are likely hitting the expected target, we asked whether a strain carrying a weak loss-of-function allele of *plk-1* is hypersensitive to the compounds. We found that a strain carrying a temperature-sensitive allele of *plk-1(or683)* exhibits 2-fold hypersensitivity to the 2,4-Dianilinopyrimidine GSK1520489 compared to the wild type control strain at the intermediate temperature of 20°C (Figure 9). The *plk-1(or683)* mutant showed an even greater sensitivity to the BT inhibitors, up to 32-fold relative to the wild type control. While not conclusive, this data is consistent with the hypothesis that the BT core scaffold is acting on the expected PLK-1 target in nematodes. Of those compounds available for retesting, the structurally related BT analogs GSK580432 and GSK479719 are the most potent across the species tested (Figure 9) and may serve as valuable structures on which to base the generation of nematode-specific PLK-1 inhibitors.

### Considering Essential Nematode-Specific Kinases as Anthelmintic Targets

An obvious alternative approach to the one we have taken here is to pursue essential nematode-specific kinases as anthelmintic targets. As described by Manning [34], there are 15 nematode-specific kinase families within the *C. elegans* kinome, consisting of 105 nematode-specific kinase genes. Of these, disruptions of 16 lead to lethal, sterile, embryonic lethal or larval arrest phenotypes (WormBase, Supplementary Table 2). These nematode-specific essential kinases belong to families within the largely expanded CK1 group, the FGFR-like RTK family KIN-16 and three worm-specific families from the Other group, which bear little resemblance to any vertebrate kinases [34]. Outside of the nematode-specific families, we identified 10 additional nematode-specific kinases with essential functions from families identified by Manning as expanded in nematodes [34]. These kinases primarily emerged from the expansion of the Fer kinase family in the worm [34].

Of these 26 essential nematode-specific kinases, we find that 23 are more than 30% identical and/or align with over 70% of the kinase domain of the closest human kinase (Supplementary Table 2). The remaining three kinases K09E4.1, F54F7.5 (MES-1), and K09C6.8 do not have detectable orthologs encoded within many of the parasitic nematodes surveyed (Supplementary Figure 4 K-M). We conclude that targeting nematode-specific kinases with the goal of developing a reasonably broad-spectrum anthelmintic is not a simple solution.

## Discussion

Here, we have taken a small molecule screening approach to survey the *C. elegans* kinome and have identified 17 druggable essential nematode kinases (Table 1). In doing so, we have identified a suite of compounds that likely inhibit these kinases *in vivo* (Supplementary Table 1). Of these 17 druggable kinases, three (EGFR/LET-23, MEK1/MEK-2 and PLK1/PLK-1) are candidate anthelmintic targets because they harbor distinct amino acid residues in the presumptive drug-binding pocket when compared to the vertebrate ortholog (Table 1). These differences may be exploited to design small molecules that specifically and safely inhibit the essential nematode kinase within the context of an infected vertebrate host.

Our strategy to identify candidate anthelmintic targets began with identifying vertebrate kinase inhibitors that are active in whole-animal *C. elegans* assays. This allowed us to uncover both the nematode kinases that can be pharmacologically manipulated *in vivo* to produce deleterious effects as well as to identify the associated small molecule scaffolds that can reach their target in whole worms. These chemical scaffolds can be refined to increase specificity for the nematode ortholog of their target kinase.

There are many advantages to using the free-living *C. elegans* nematode as a platform to identify small molecules with anthelmintic potential. First, it is cost-effective and can be adapted to medium or high-throughput screens. Second, molecules found to be active against *C. elegans* have an increased likelihood of having activity against parasitic nematodes [66, 67]. Third, *C. elegans* small molecule bioassays have proven to be exceptionally useful in identifying and/or characterizing several candidate anthelmintics [66, 68-72]. Fourth, standard 6-day *C. elegans* bioassays of the type we have employed here probe the entire life cycle of the nematode. Hence, this approach can reveal nematode vulnerabilities that may not be accessible using bioassays with parasitic species, which are often limited to a single life stage. Finally, the wealth of genetic knowledge and *C. elegans* tools available allow us to make inferences about target engagement of bioactive molecules based on established genetic loss-of-function phenotypes, and can further help validate the target of these molecules with relative ease (this work) [66, 69, 71, 72].

Here, we have used *C. elegans* to demonstrate the anthelmintic potential of EGFR/LET-23, MEK1/MEK-2 and PLK1/PLK-1 kinase targets. There is corroborating evidence in the literature that supports the idea that these kinases may have value as anthelmintic targets. The EGFR/ERK pathway has been suggested as a therapeutic target in the tapeworm *Echinococcus multilocularis*, the causative parasite of alveolar echinococcosis [16, 73]. In *E. multilocularis*, the EGFR/ERK signaling pathway has been implicated in promoting proliferation of the germinative cells, a stem cell-like population that drives larval growth and development. Exposure of *E. multilocularis* to the EGFR inhibitors CI-1033 and BIBW2992, or the MEK/ERK inhibitor U0126, impairs germinative cell proliferation, resulting in larval growth and development defects [16]. Furthermore, the *E. multilocularis* PLK1 homolog EmPlk1 is also expressed in the germinative cells and can be inactivated by the PLK1 inhibitor BI2536, inducing mitotic arrest and germinative cell death [74].

Other studies have implicated the EGFR/ERK signaling pathway in the flatworm parasite *Schistasoma mansoni* as a target to treat schistosomiasis due to its role in the development of oocytes and the female somatic gonad [75]. A screen of the 114 compounds in the National Cancer Institute’s Oncology Drug Set identified 11 compounds that had an effect on both *S. mansoni* adults and larvae *in vitro* with an IC_50_ ≤ 10 µM [76]. These included the MEK1/2 inhibitor trametinib and three tyrosine kinase inhibitors annotated to target EGFR (gefitinib, afatinib and vandetanib). Trametinib and vandetanib maintained activity *in vivo* and were found to reduce worm burden by 63.8% and 48.1% respectively in the chronic *S. mansoni* mouse model of infection after a single oral dose of 400 mg/kg [76]. Hence, nematode-specific EGFR and MEK-2 inhibitors may have broad-spectrum utility.

PLK1 has also been investigated as an anthelmintic target in *S. mansoni* [21]. RNAi*-*knockdown of the *S. mansoni* PLK1 ortholog smPLK1 had a deleterious effect on post-infective larvae (schistosomulae). Small molecule inhibitors of human PLK1 were found to induce uncoordinated movements in adults and morphological defects in both schistosomulae and adult parasites. The benzimidazole thiophene core scaffold of PLK1 inhibitors from the GSK PKIS shows bioactivity in *S. mansoni* and was identified as a potential anti-schistosomal scaffold [21]. In the work presented here, we have also identified the benzimidazole thiophenes as a nematicidal scaffold that likely inhibits worm PLK-1, further reinforcing the idea that PLK-1 may be a useful anthelmintic target (Figure 9; Supplementary File S2D).

A number of the kinase inhibitors that we found to have robust activity in *C. elegans* target vertebrate proteins that have no obvious ortholog in *C. elegans* (Supplementary Table 1). These hits include the 9 unique inhibitors that target vertebrate RET kinase and 17 unique inhibitors that target the vertebrate PDGFR family members PDGFRA, PDGFRB, KIT, FLT3 and CSF1R. The targets of other active inhibitors have a *C. elegans* ortholog, but these orthologs lack essential functions in *C. elegans* (Supplementary Table 1). For example, there are 16 unique compounds that inhibit mammalian VEGFR receptor family members FLT1, KDR and FLT4 that we found to elicit dramatic phenotypes in *C. elegans*. However, the corresponding *C. elegans* orthologs, VER-1, VER-3 and VER-4, have no reported phenotypes on WormBase, despite the characterization of both their genetic loss-of-function and RNAi phenotypes. The phenotypes elicited by these 16 compounds may be the result of promiscuous inhibition of multiple kinases or due to off-target effects. Deciphering the targets of these lethal compounds in *C. elegans* would be a valuable next step to uncover additional anthelmintic targets.

One alternative approach is to focus on developing inhibitors against nematode-specific kinases with no structurally similar match in vertebrate host species, such as the three we highlight here (K09E4.1, F54F7.5/MES-1, and K09C6.8, Supplementary Table 2). There are two clear challenges associated with this alternative. First, identifying inhibitors that specifically target the nematode-specific kinases and do not inhibit a wide array of kinases (including those in the host) is a formidable challenge. In the approach that we have focused on here, extensive work has already gone into optimizing structures to specifically target EGFR, MEK1/MEK2, and PLK1. Second, and perhaps more importantly, nematode-specific kinases may be more evolutionarily divergent within nematoda than those conserved with vertebrates (Supplementary Figure 4) and it is unknown if the essential *C. elegans* function of any of these kinases is conserved with parasitic nematode species. Hence, any inhibitor that targets a given nematode-specific kinase may have limited utility against other nematode parasites.

Ample evidence shows that anthelmintic resistance is rampant among parasitic nematodes infecting livestock [13], and resistance among human parasites is growing [2]. There is a clear need for the development of new anthelmintic drugs to add to our arsenal. Here, we have highlighted three essential kinases that have good anthelmintic potential because of small sequence changes in the drug-binding pockets of the nematode orthologs relative to mammalian hosts. These kinases have proven to be druggable in whole worms, making them important targets for the development of novel anthelmintic drugs.

## Methods

### Nematode Strains and Culture Methods

*Caenorhabditis elegans* N2 (wild-type), CB1275 *lin-1(e1275), C. briggsae* AF16 (wild-type) and *Pristionchus pacificus* PS312 (wild-type) were provided by the *Caenorhabditis* Genetics Center (University of Minnesota). The JA337 strain *let-23(sy1);* jgIs19[let-23p::LET-23::hEGFR-TK, rol-6(su1006)] was provided by Dr. Jaegal Shim. All animals were cultured using standard methods at 20°C [40], with the exception of the temperature sensitive mutant *plk-1(or683)* which was cultured at 15 °C.

### Chemical Sources

The Library of Pharmacologically Active Compounds (Sigma-Aldrich) was purchased from the SickKids Proteomics, Analytics, Robotics & Chemical Biology Centre (SPARC BioCentre). The APExBIO DiscoveryProbe Kinase Inhibitor Library was a gift from Jim Dowling. The OICR Kinase Inhibitor Library was a gift from Rima Al-Awar and David Uehling at the Ontario Institute of Cancer Research. The GSK Published Kinase Inhibitor Set (PKIS) molecules was obtained from GlaxoSmithKline. MEK Inhibitors trametinib, pimasertib, U0126-EtOH and BI-847325 and EGFR inhibitor gefitinib were purchased for further testing from Selleck Chemicals.

### Kinase Inhibitor Screens

The kinase inhibitor screening method was adapted from previously described liquid-based screening protocols [66] and is summarized in Figure 1A. Briefly, *Escherichia coli* strain HB101 was resuspended in nematode growth media (NGM) buffered using Potassium Phosphate buffer (pH=6) [77] or Citrate Phosphate buffer (pH=3); [78]. The final media pH for these two conditions were pH=7 (“neutral” media) and pH=4.5 (“acidic” media) respectively. The acidic media condition was included to improve our ability to capture phenotypes induced by molecules that are charged at neutral pH and thus may not be bioavailable to the worms. 40 μL of bacterial suspension was dispensed into each well of a 96-well plate, and 300 nL of the small molecule inhibitors or of dimethyl sulfoxide (DMSO) vehicle control was pinned into the wells using a 96-well pinning tool (V&P Scientific). Approximately 20 synchronized first-stage larvae (L1s) N2 worms obtained from an embryo preparation were then added to each well in 10 μL M9 buffer (neutral media) or NGM (acidic media) [77]. The final concentration of DMSO in the wells was 0.6% v/v. Plates were sealed with Parafilm and incubated for 6 days at 25°C while shaking at 200 rpm (New Brunswick I26/I26R shaker, Eppendorf). On day 6 the plates were observed under a dissection microscope and embryonic lethality, sterility, larval arrest and lethality phenotypes were assessed in each condition.

The LOPAC library was screened once in its entirety in technical duplicate in both media conditions. Kinase inhibitor hits in either media condition identified in the primary screen were retested in duplicate in both media conditions. The OICR library was screened twice in its entirety in technical duplicate in both media conditions. The APExBIO library was screened in technical duplicate in neutral pH media only due to limited drug availability. The PKIS library was screened once in its entirety in technical duplicate in both media conditions. Hits in either media condition identified in the primary screen were retested in duplicate in both media conditions based on availability.

Hit compounds were defined by the following criteria: in the primary screens, compounds were considered hits if the inhibitor induced population growth defects by day 6 (both technical replicates have fewer than 50 worms, at least one of the two replicates has fewer than 25 worms) and/or relevant phenotypes of interest (Emb, Ste, Lva, Let, Bag, Rod-like progeny) in both technical replicates in either media condition. Inhibitors were included in the Supplementary File S1A ‘Hit Summary’ and considered hits for further analysis if: 1) upon retest, the inhibitor induced population growth (both technical replicates have fewer than 50 worms, at least one of the two replicates has fewer than 25 worms) and/or relevant phenotypes of interest (Emb, Bag, Ste, Lva, Let, Rod-like progeny) in both technical replicates as in the primary screen; 2) the inhibitor was considered a hit in the primary screen and we were not able to repeat due to insufficient molecule.

Follow up inhibitor dose response analyses were performed using the same methodology described above using neutral media unless otherwise specified.

### Kinase Inhibitor Target Assignment

For the ‘commercial’ kinase inhibitor libraries (LOPAC, OICR and APExBIO) the library documentation included annotations for the kinases targeted by each inhibitor within the libraries. For the PKIS library, we used the published *in vitro* kinase inhibition data to assign kinase targets to each inhibitor hit [36-38]. For hits from PKIS1, any kinase that was inhibited ≥65% at 0.1 μM compound concentration was considered a target. For hits from PKIS2, any kinase that was inhibited ≥90% at 1 μM compound concentration was considered a target. If there were more than 10 kinase targets for a particular PKIS inhibitor according to these criteria, the inhibitor was annotated as having ‘MANY’ targets and the specific targets of these compounds were not included in the downstream analyses.

### Homology Modeling and Sequence Analysis

*C. elegans* orthologs of human kinases targeted by our inhibitors were identified using OrthoList 2 [79]. *C. elegans* kinase protein structure homology models were generated using the SWISS-MODEL pipeline using the indicated structural templates accessed from the Protein Data Bank (PDB) [42]. At least two unique human kinase templates were used for modeling each nematode kinase ortholog if available. The structures were visualized using The PyMOL Molecular Graphic System (Version 2.1.1 Schrödinger, LLC). All residues lining the inhibitor binding pocket(s) for each kinase of interest were analyzed to identify divergent AA residues with side chains oriented towards the inhibitor binding site. Particular attention was given to those residues within close proximity to the bound inhibitor molecule (within 5Å as measured using PyMOL).

Multiple sequence alignments of the kinase domains across vertebrate and nematode species were generated using Clustal Omega [80]. Relevant protein sequences from the following nematode and vertebrate species were identified using NCBI BLAST for inclusion in the alignments: free-living nematodes *C. elegans* (Clade V), *C. briggsae* (Clade V), *P. pacificus* (Clade V); parasitic nematodes *Haemonchus contortus* (Clade V), *Necator americanus* (Clade V), *Ancylostoma duodenale* (Clade V), *Dictyocaulus viviparus* (Clade V), *Ancylostoma ceylanicum* (Clade V), *Strongyloides ratti* (Clade IV), *Brugia malayi* (Clade III), *Onchocerca ochengi* (Clade III), *Loa loa* (Clade III), *Trichuris trichiura* or *T. suis* (Clade I); vertebrates *Danio rerio, Gallus gallus, Mus musculus, Rattus norvegicus, Canis lupus familiaris, Ovis aries, Bos taurus* and *Homo sapiens*.

### Vulva Phenotype Analysis

Inhibitor exposure was performed in liquid media as described in the Liquid-Based Kinase Inhibitor Screening method above. Analysis was performed once all F0 worms reached adulthood (Day 4). For the MEK inhibitor experiments, worms were mounted on a 2% agarose pad on a glass slide, observed using a 20x objective on a Leica DMRA microscope and the number of vulval protrusions were counted. For the EGFR inhibitor experiment, the adult worms were examined under a dissection scope and the vulva phenotype was categorized as Multivulva, WT or Vulvaless. F0 worms that contained hatched progeny trapped inside (bag-of-worms phenotype) were assumed to not have a functioning vulva and were counted as Vulvaless. A minimum of three biological replicates were completed for all vulva phenotype analyses.

## Figure Legends

Supplementary Table 1. Kinases Targeted by Inhibitor Screen Hits

Supplementary Table 2. Essential Nematode-Specific Kinases

Supplementary File S1. *C. elegans* Vertebrate Kinase Inhibitor Screen Results.

S1A: Summary of 191 hits from kinase inhibitor screen

S1B: Full screen results for LOPAC library kinase inhibitors

S1C: Full screen results for OICR kinase inhibitor library

S1D: Full screen results for APExBIO Discovery Probe kinase inhibitor library

S1E: Full screen results for PKIS1 and PKIS2

Supplementary File S2. EGFR and PLK1 inhibitors included in screening set.

S2A: Commercial EGFR inhibitor screening results

S2B: Screening results for PKIS inhibitor scaffolds targeting EGFR

S2C: Commercial PLK1 inhibitor screening results

S2D: Screening results for PKIS inhibitor scaffolds targeting PLK1

---

**Supplementary Figure 1.**
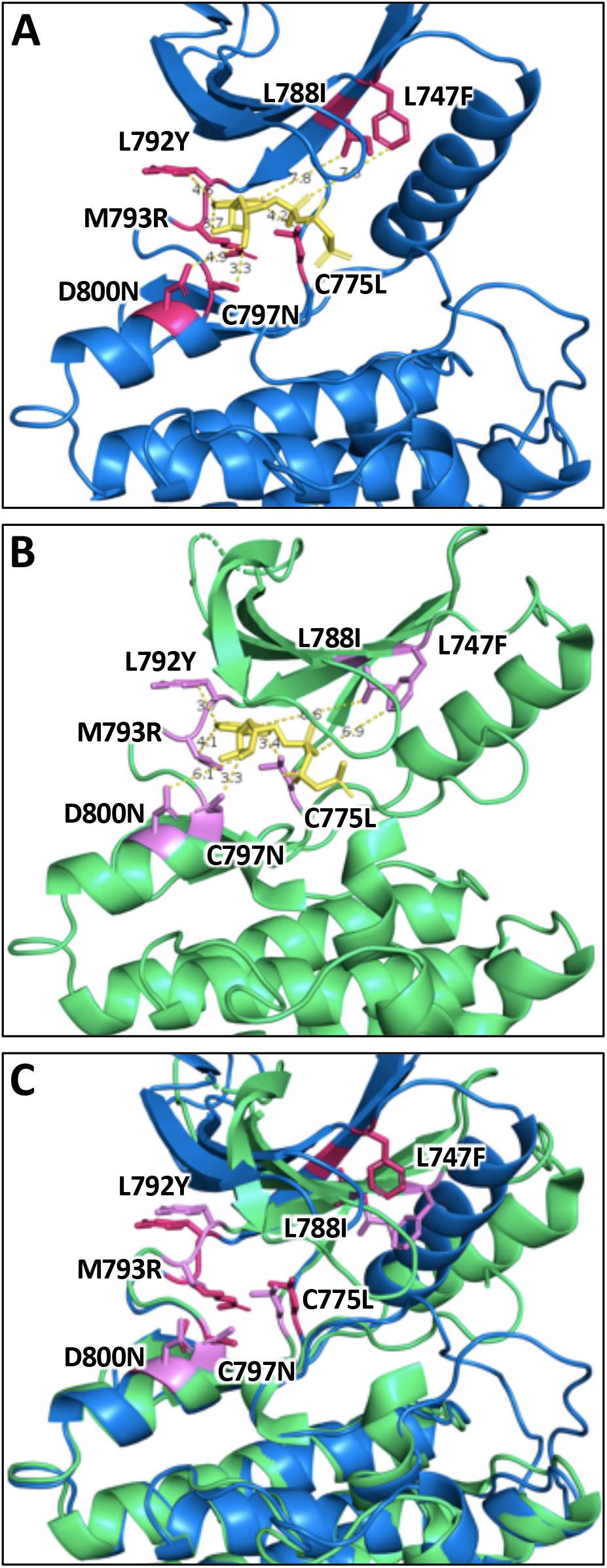
Structural comparison of EGFR/LET-23 homology model and LET-23 crystal structure. (A) Homology model of *C. elegans* LET-23 modeled after an EGFR crystal structure complexed with AMP-PNP (PDB: 2ITX, AMP-PNP in yellow). The numerous divergent residues between *C. elegans* LET-23 and Human EGFR with side chains oriented towards the active site are highlighted in pink. Residues are labeled according to position in human EGFR, with the first letter indicating the identity of the human residue and the latter indicating the identity of the *C. elegans* residue. (B) The crystal structure of the LET-23 kinase domain complexed with AMP-PNP (PDB: 5WNO, AMP-PNP in yellow) with the same residues highlighted in purple as in (A). The orientation of these key residues towards the active site is consistent between the LET-23 homology model and crystal structure. The homology model and crystal structure of LET-23 are overlaid in (C).

**Supplementary Figure 2.**
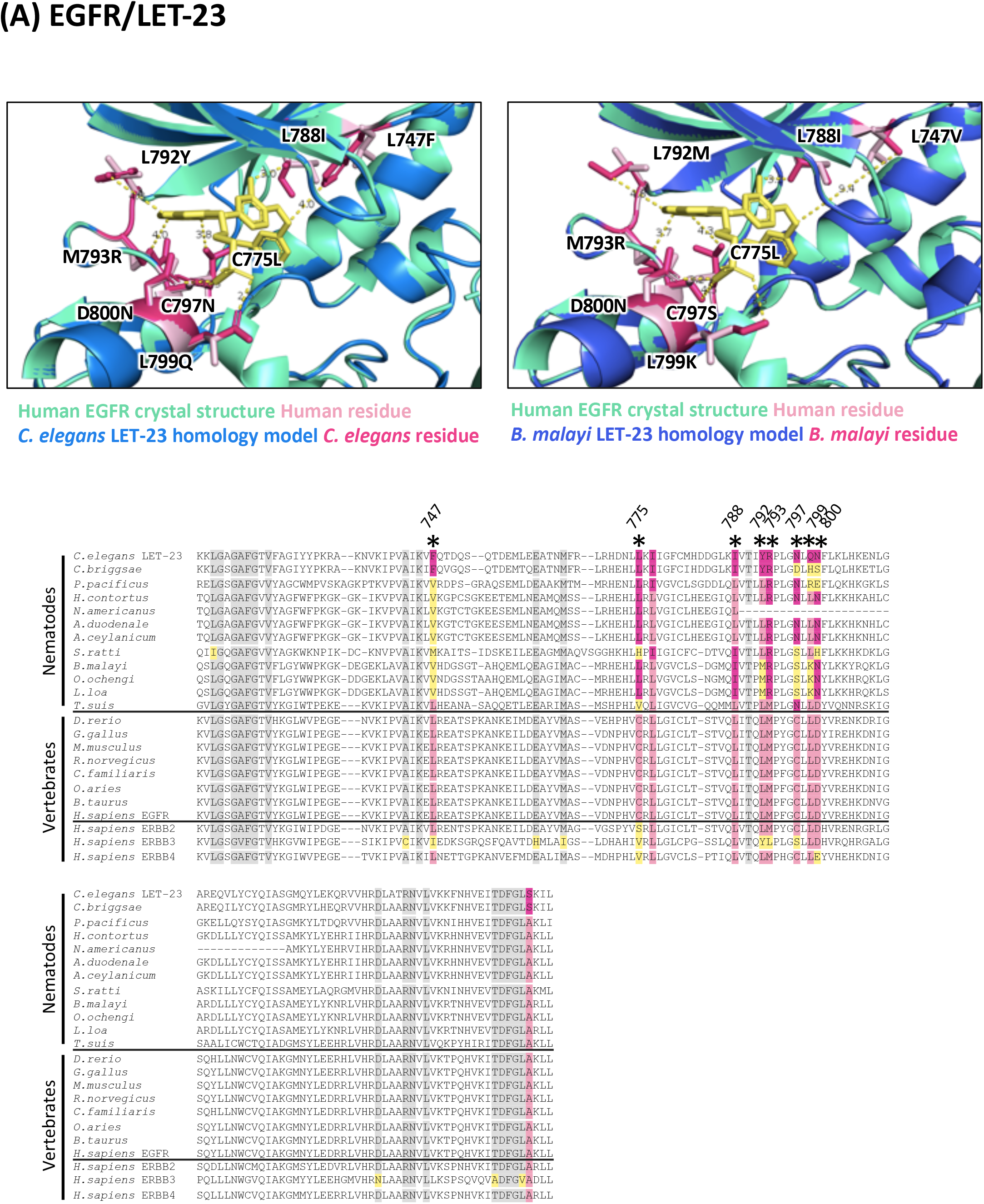
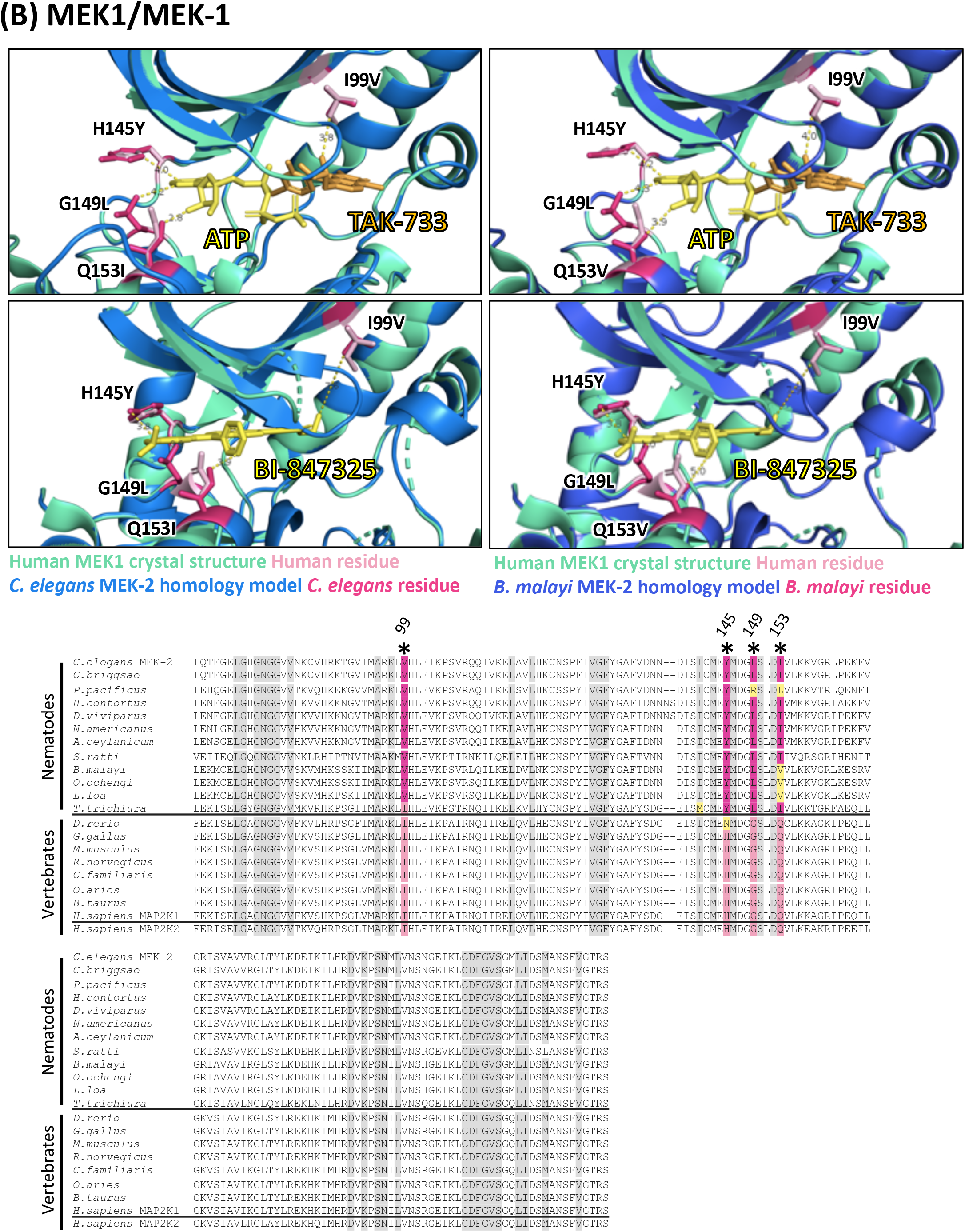
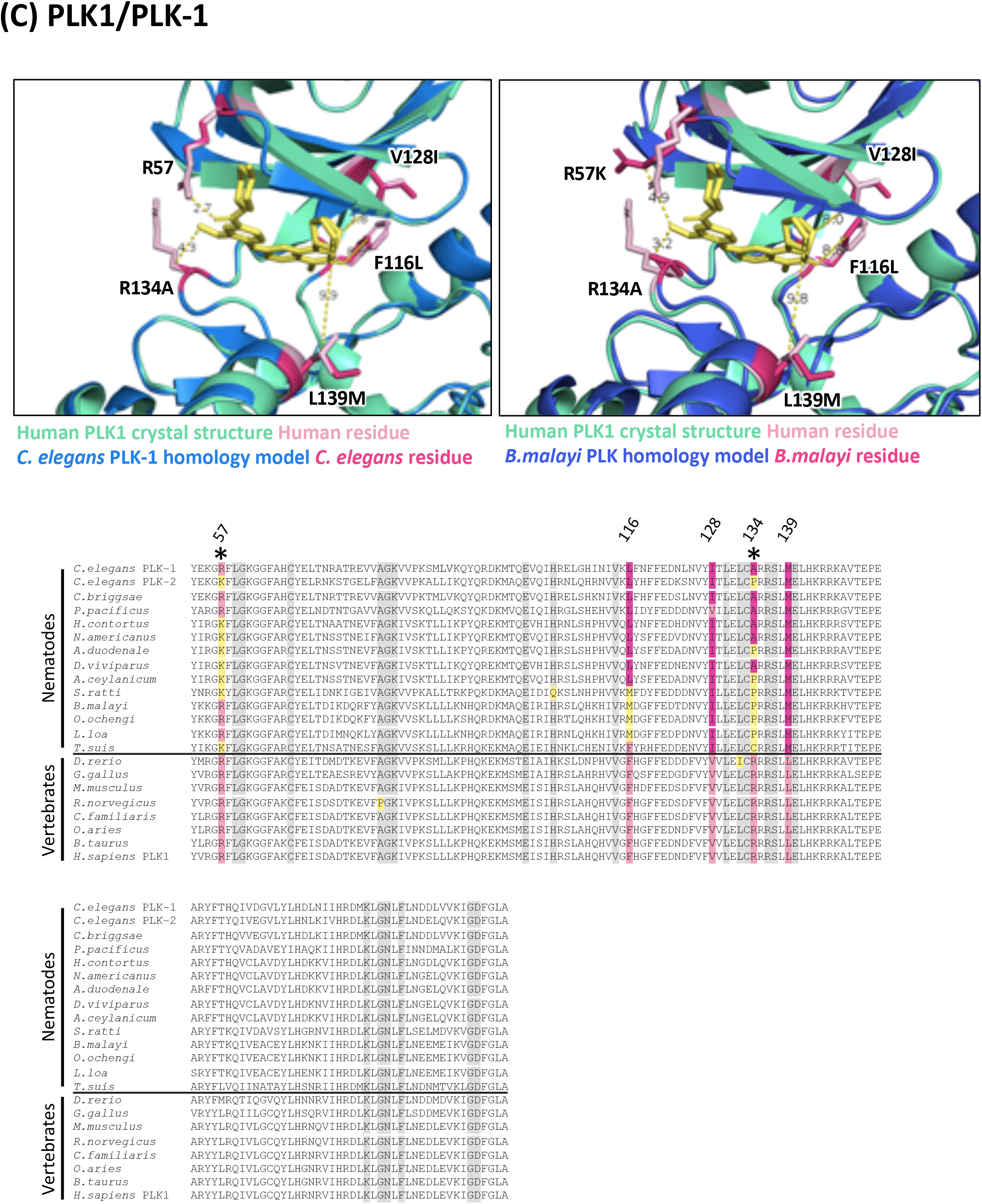
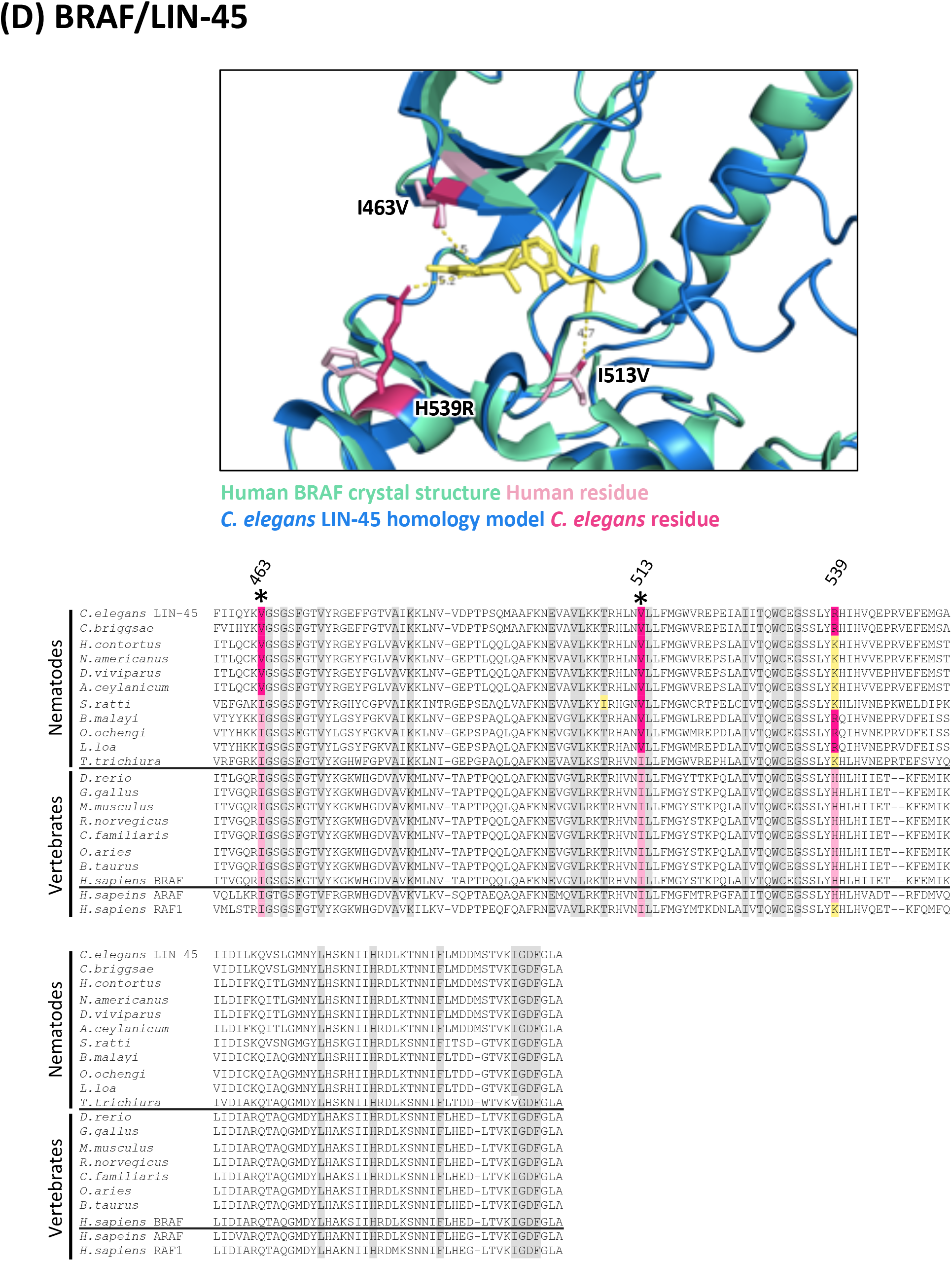
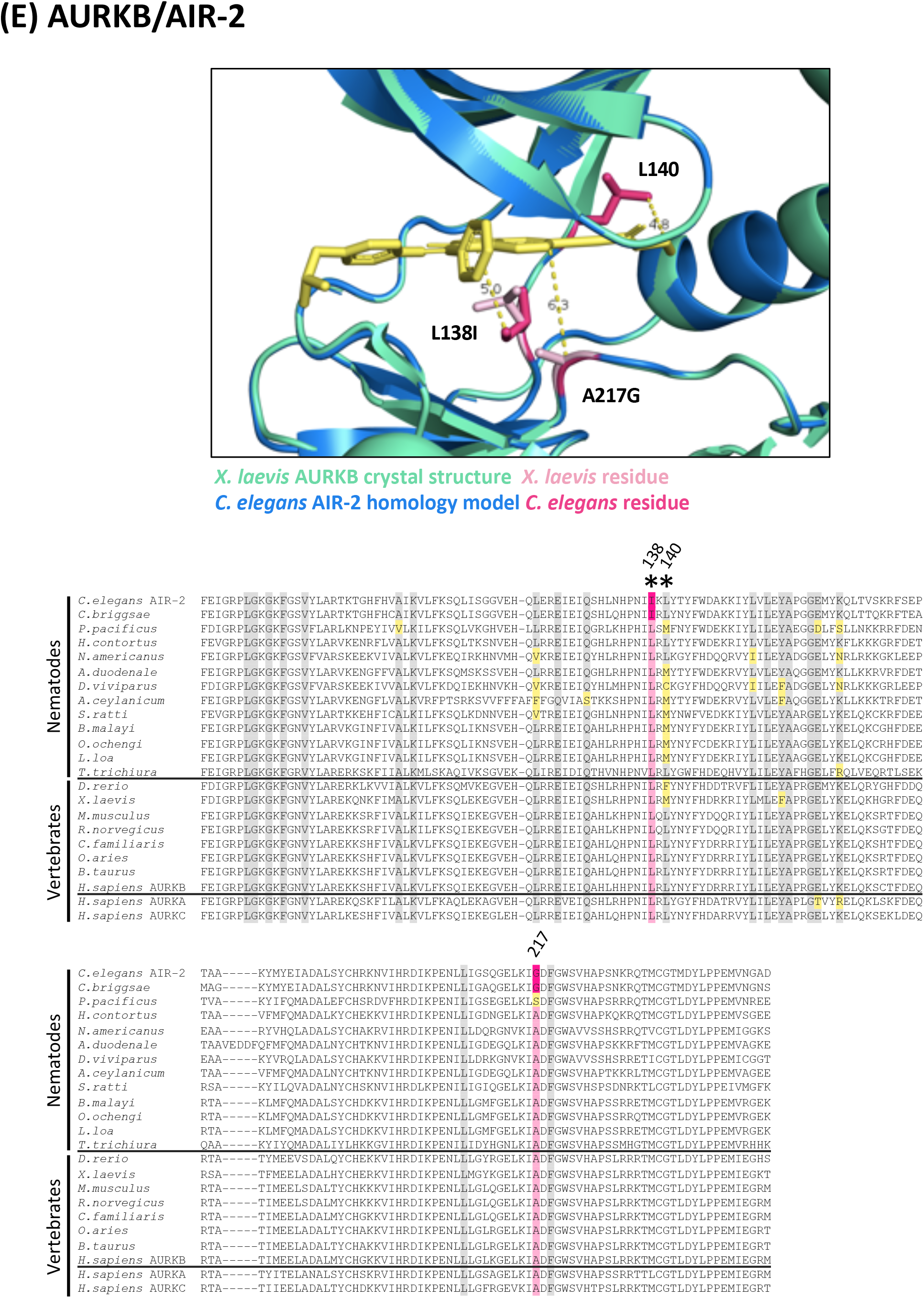
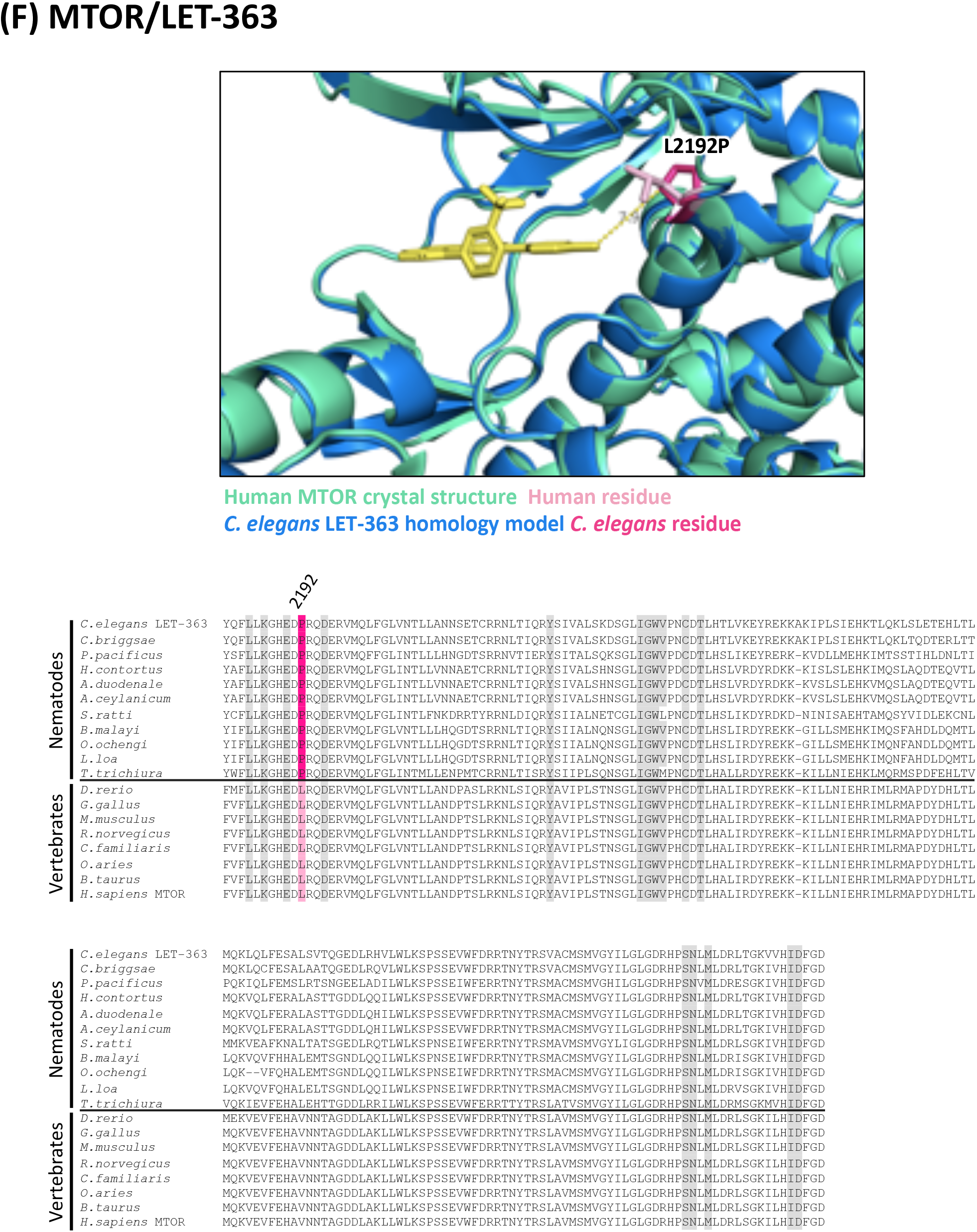
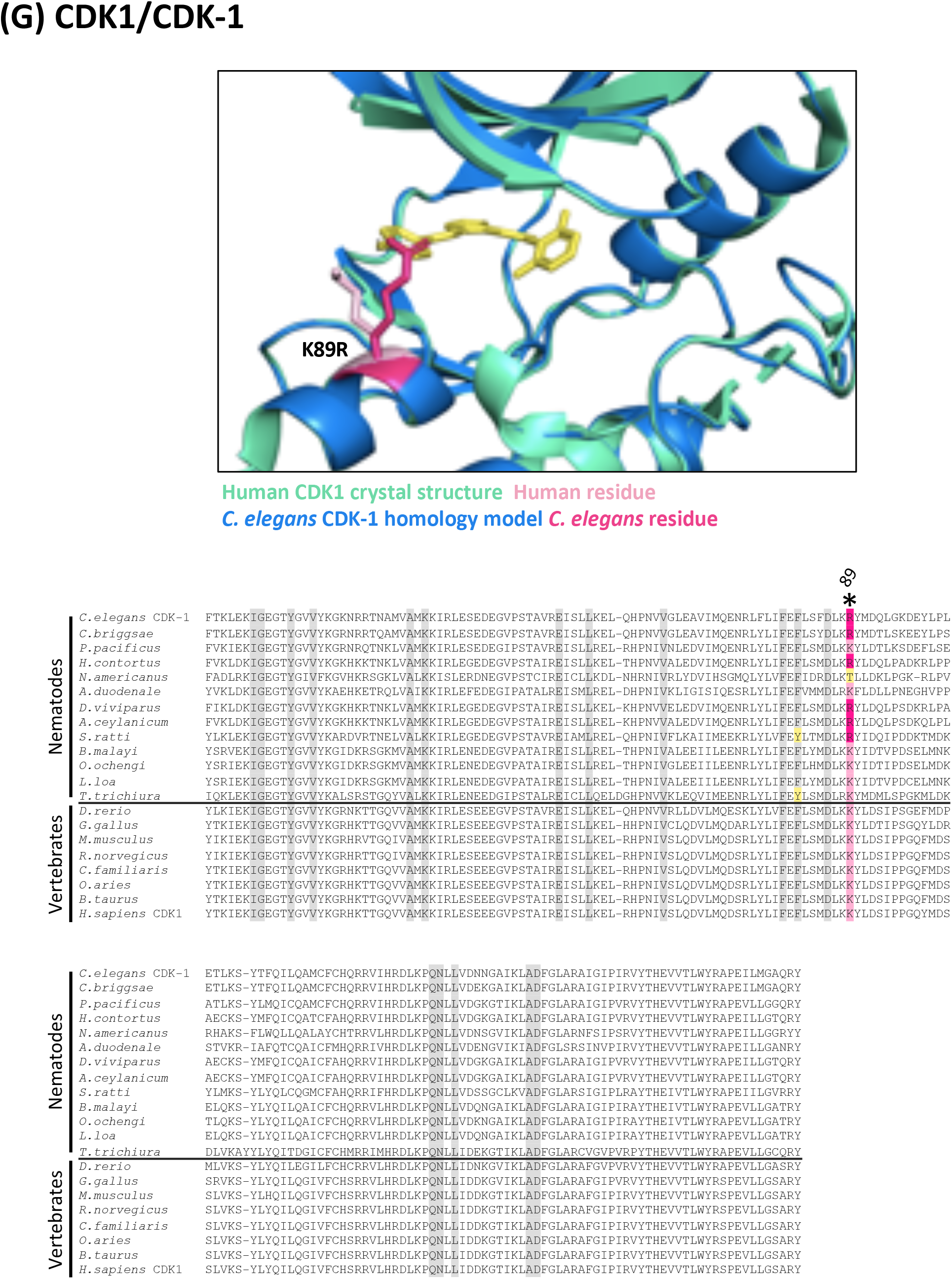
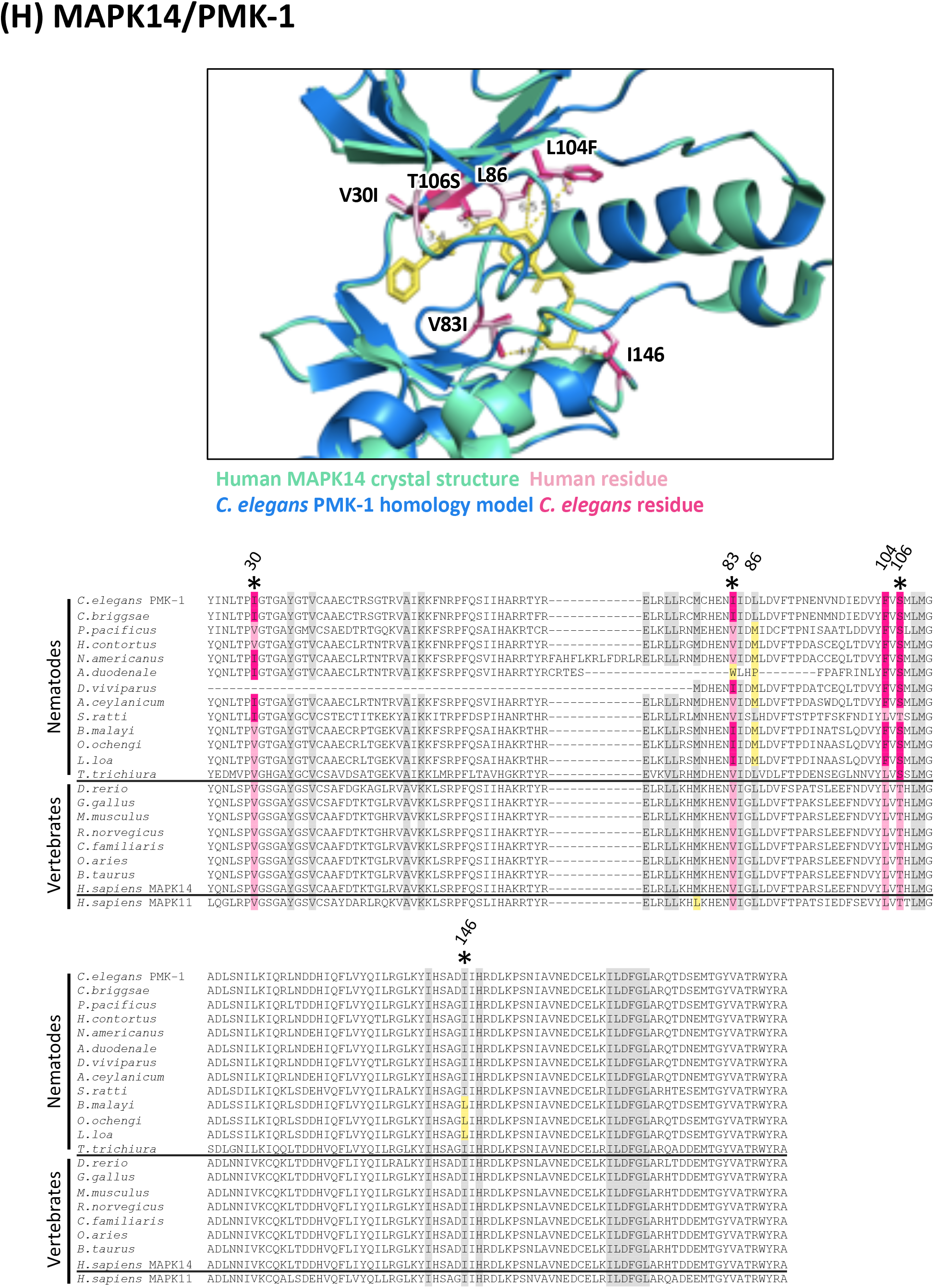
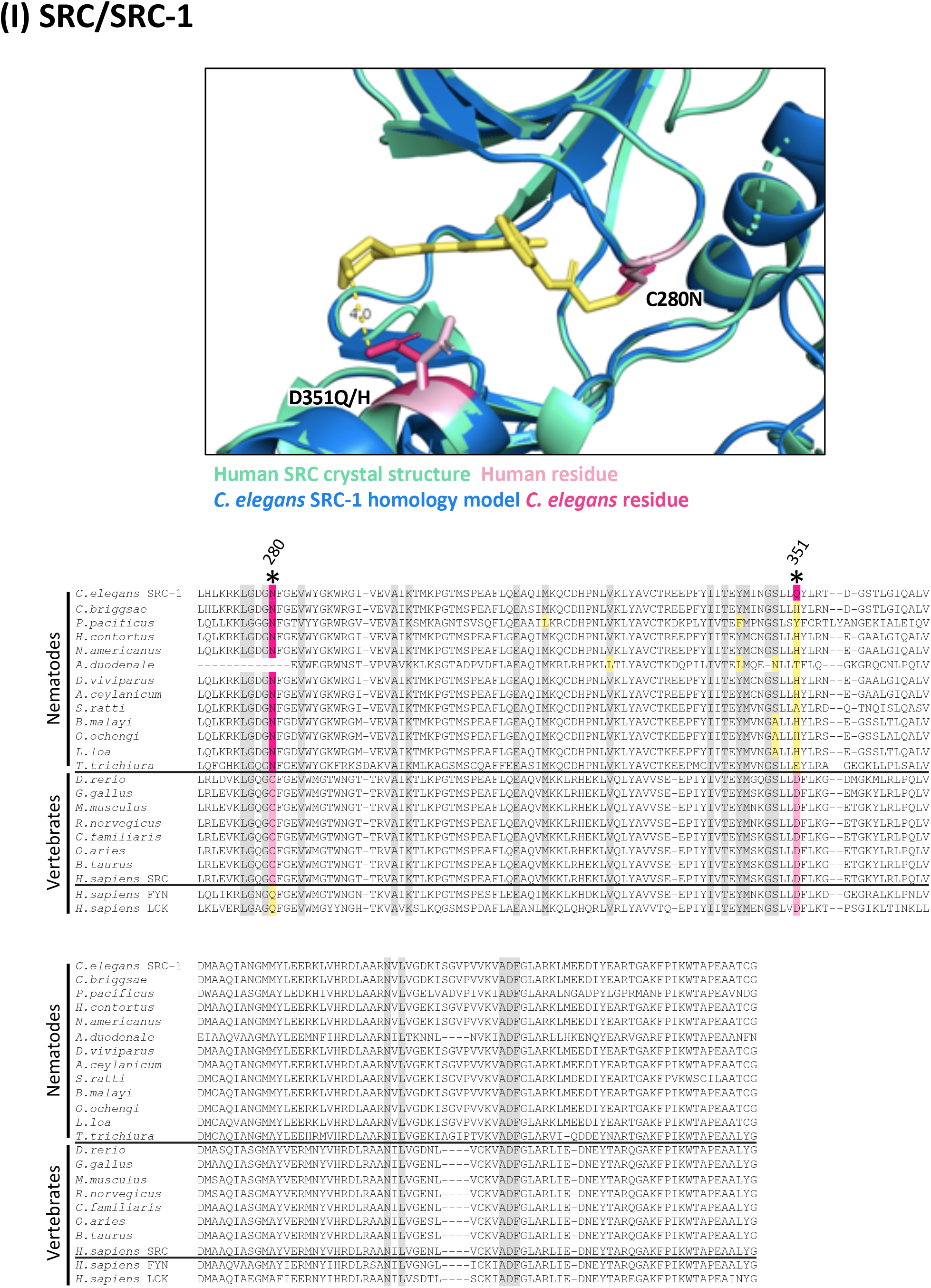
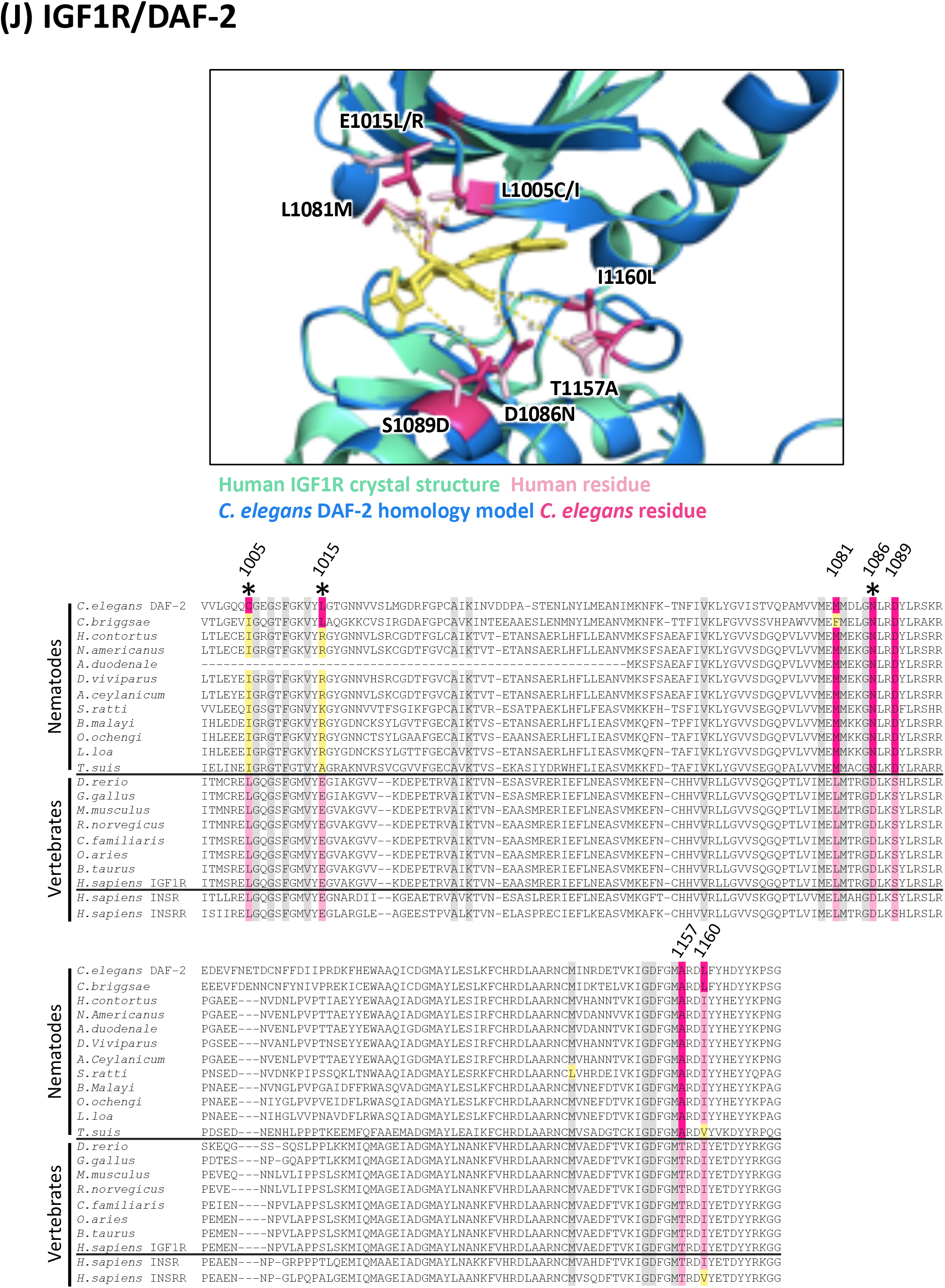
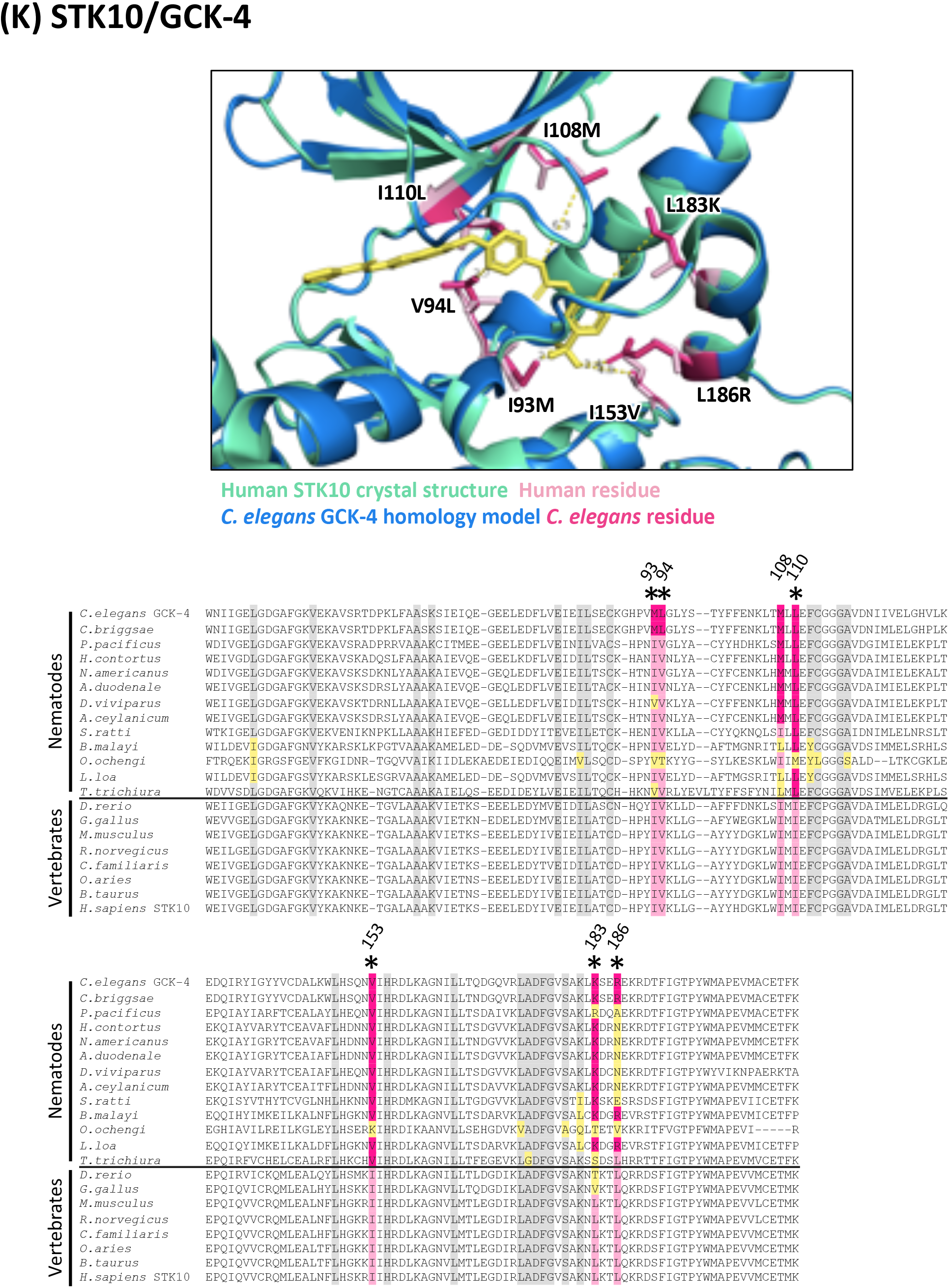
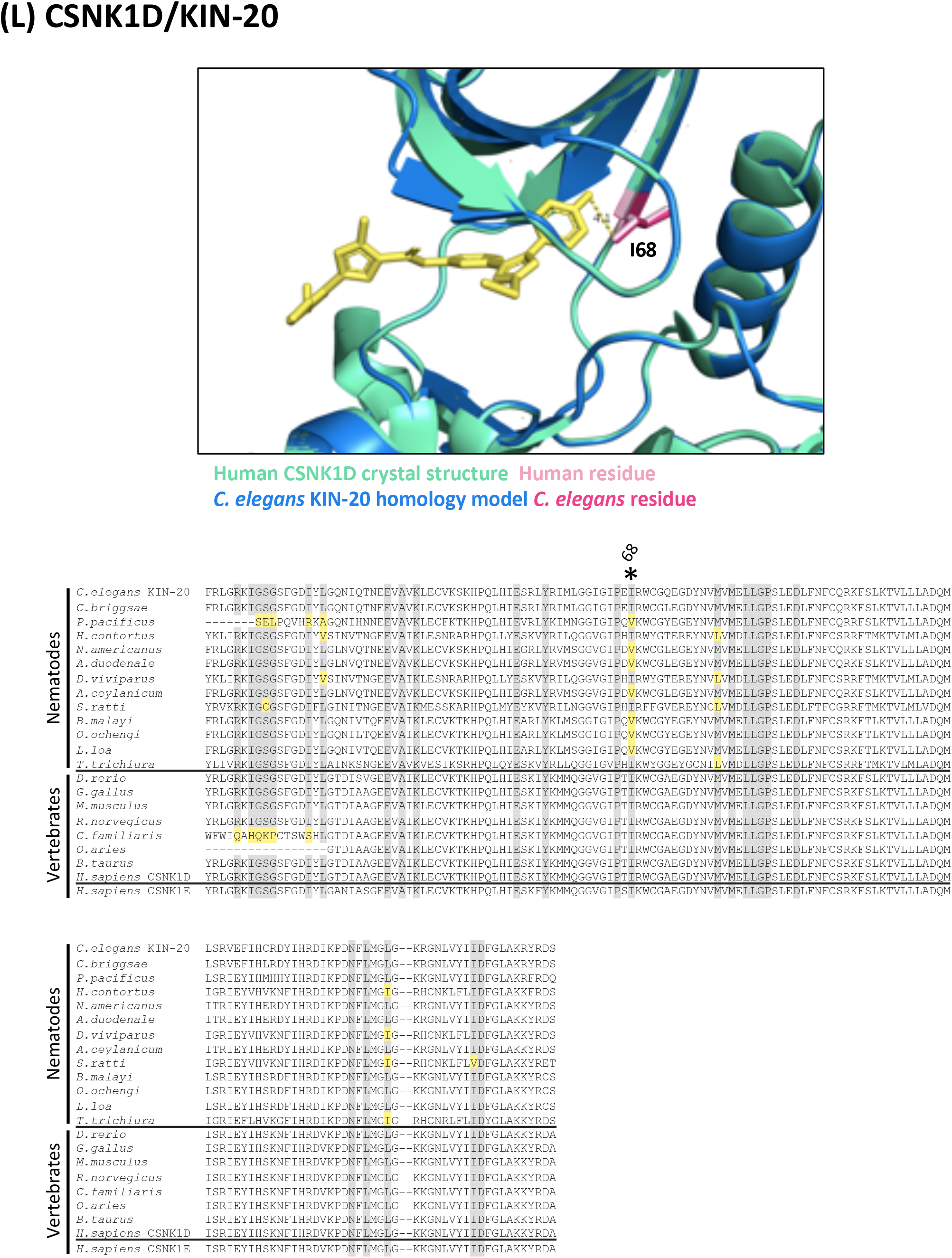
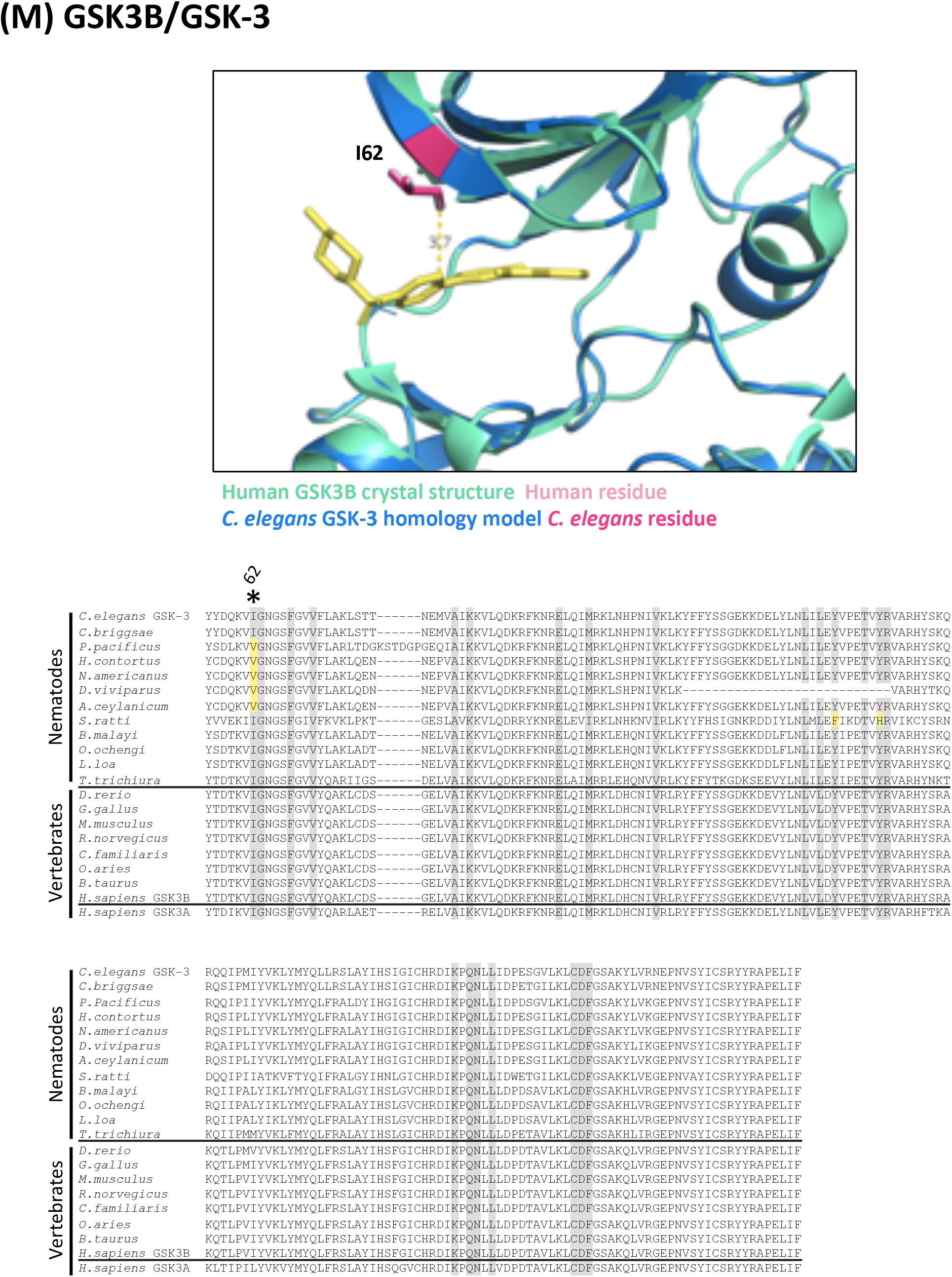
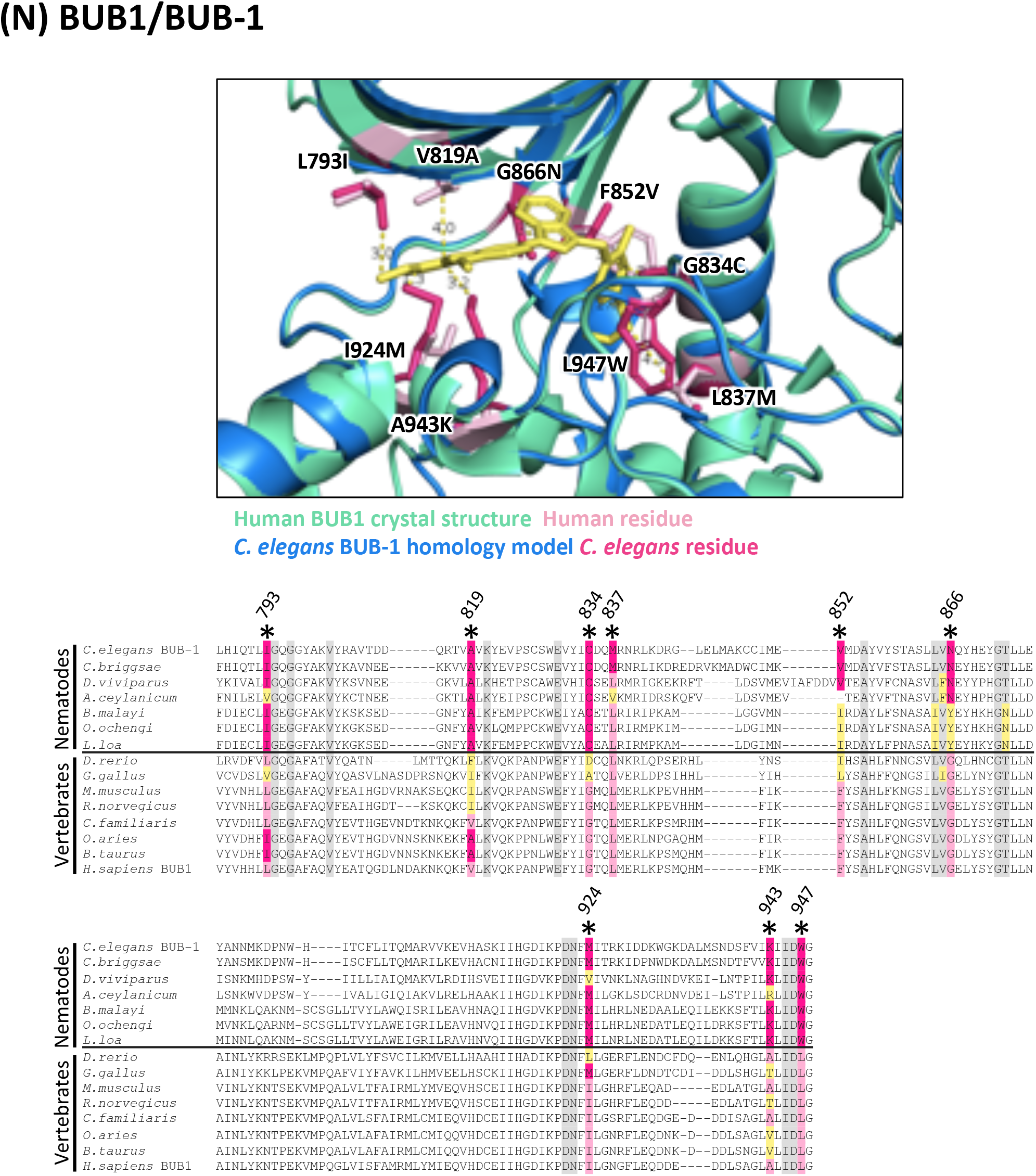
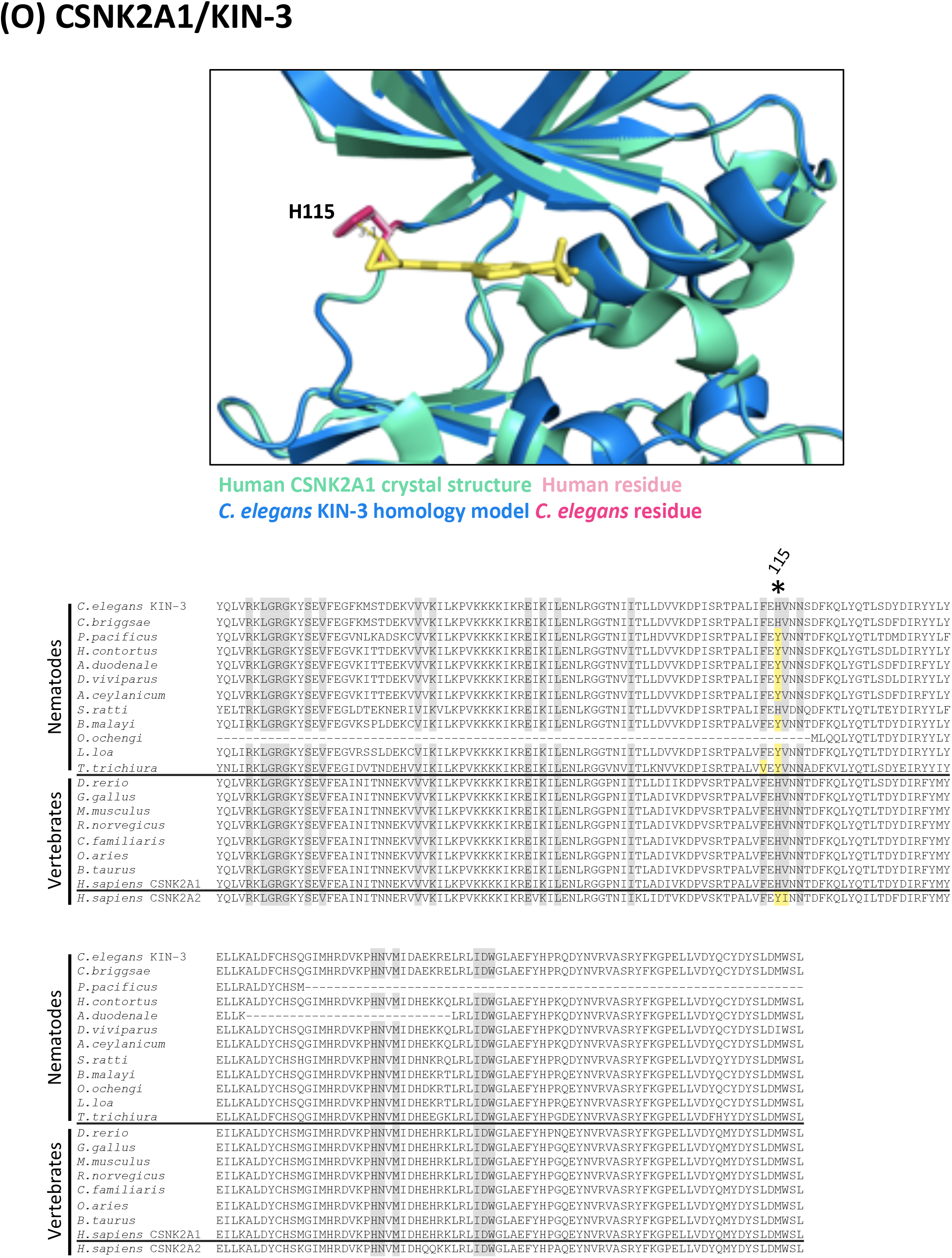
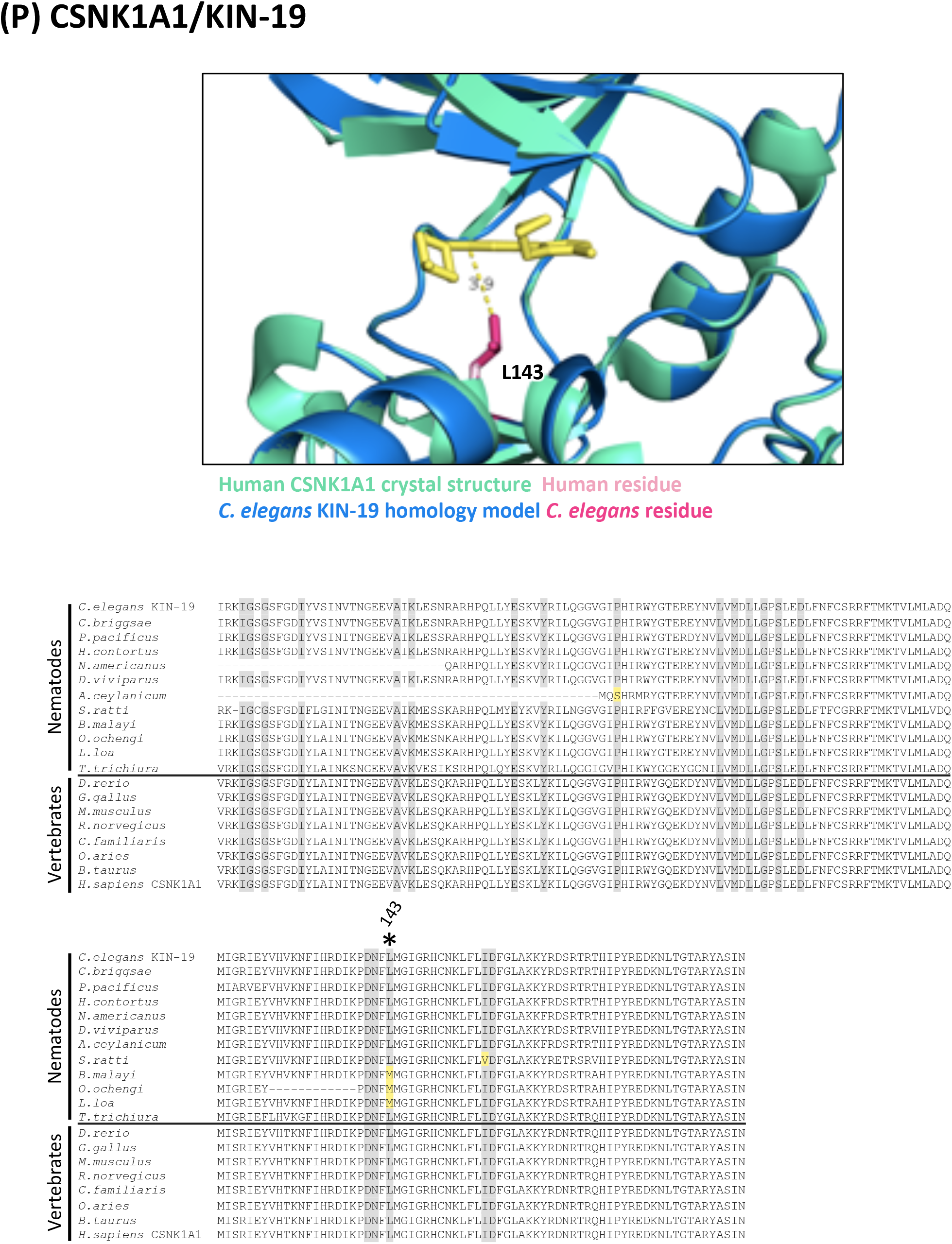
Structures and alignments for druggable nematode essential kinase targets. In all alignments, residues that are proximal to the inhibitor binding site with side-chains facing inwards are highlighted on the alignment. Grey residues were examined but do not differ between *C. elegans* and humans. Divergent residues are highlighted in dark pink *(C. elegans* residue) and light pink (human residue). Yellow highlights those residues that differ in identity from both human and *C. elegans* in these key positions. Residues of interest that differ between nematodes and vertebrates are labeled with their position in the human kinase on the alignment and shown in pink on the structure above. Those residues of interest indicated with a star on the alignment are located within 5Å of the bound inhibitor in any of the *C. elegans* models examined. **(A) Structure and alignment of EGFR and LET-23**. Structure of PDB:1XKK (Human EGFR) in green with Lapatinib bound (in yellow), aligned to *C. elegans* LET-23 homology model in blue. The human and *C. elegans* kinase domains share 44% identity. All residues that are proximal to the inhibitor binding site with side-chains facing inwards are highlighted on the alignment (identified from PDB structures 1XKK, 2ITX, 3W2S, 4G5J and associated *C. elegans* homology models). The orthologous LET-23 sequence from the parasitic nematode *B. malayi* was modeled to the human EGFR structure (PDB: 1XKK) to confirm the favourable position of the identified divergent residues within the inhibitor binding site. Residues of interest are indicated on the structures in dark pink (nematode residue) and light pink (human residue). **(B) Structure and alignment of MEK1 and MEK-2**. Structure of PDB: 3PP1 (Human MEK1) in green with allosteric inhibitor TAK-733 bound (in orange) aligned to *C. elegans* MEK-2 homology model in blue above. Structure of PBD:5EYM (Human MEK1) with ATP-competitive inhibitor BI-847325 bound (in yellow) aligned to *C. elegans* MEK-2 homology model below. The human and *C. elegans* kinase domains share 60% identity. All residues proximal to the ATP-competitive inhibitor binding site (identified in PDB structures 5EYM, 5HZE) and allosteric site (identified from PDB structures 3EQH, 3PP1) with side-chains facing inwards are highlighted on the alignment. The orthologous MEK-2 sequence from the parasitic nematode *B. malayi* was modeled to the human MEK-1 structures (PDB: 3PP1 above, PDB: 5EYM below) to confirm the favourable position of the identified divergent residues within the inhibitor binding site. Residues of interest are indicated on the structures in dark pink (nematode residue) and light pink (human residue). **(C) Structure and alignment of PLK1 and PLK-1**. Structure of PDB: 2RKU (Human PLK1) in green with BI2536 bound (in yellow), aligned to *C. elegans* PLK-1 homology model in blue. The human and *C. elegans* kinase domains share 64% identity. All residues that are proximal to the inhibitor binding site with side-chains facing inwards are highlighted on the alignment (identified from PDB structures 2RKU, 3FC2, 2YAC, 4J52 and associated *C. elegans* homology models). The orthologous PLK-1 sequence from the parasitic nematode *B. malayi* was modeled to the human PLK1 structure (PDB:2RKU) to confirm the favourable position of the identified divergent residues within the inhibitor binding site. Residues of interest are indicated on the structures in dark pink (nematode residue) and light pink (human residue). **(D) Structure and alignment of BRAF and LIN-45**. Structure of PDB: 5CSW (Human BRAF) in green with dabrafenib bound (in yellow), aligned to *C. elegans* LIN-45 homology model in blue. The human and *C. elegans* kinase domains share 62% identity. All residues that are proximal to the inhibitor binding site with side-chains facing inwards are highlighted on the alignment (identified from PDB structures 5CSW, 5CT7 and associated *C. elegans* homology models). Residues of interest are indicated on the structure above in dark pink (*C. elegans* residue) and light pink (human residue). **(E) Structure and alignment of AURKB and AIR-2**. Structure of PDB: 5EYK (*Xenopus laevis* AURKB) in green with BI-847325 bound (in yellow), aligned to *C. elegans* AIR-2 homology model in blue. The human and *C. elegans* kinase domains share 64% identity. All residues that are proximal to the inhibitor binding site with side-chains facing inwards are highlighted on the alignment (identified from human structure PDB: 4AF3 and *X. laevis* structure PDB: 5EYK and associated *C. elegans* homology models). Residues of interest are indicated on the structure above in dark pink (*C. elegans* residue) and light pink (*X. laevis* residue). **(F) Structure and alignment of MTOR and LET-363**. Structure of PDB: 4JSX (Human MTOR) in green with Torin 2 bound (in yellow), aligned to *C. elegans* LET-363 homology model in blue. The human and *C. elegans* kinase domains share 62% identity. All residues that are proximal to the inhibitor binding site with side-chains facing inwards are highlighted on the alignment (identified from PDB structures 4JSX, 4JSV and associated *C. elegans* homology models). Residues of interest are indicated on the structure above in dark pink (*C. elegans* residue) and light pink (human residue). **(G) Structure and alignment of CDK1 and CDK-1** Structure of PDB: 4Y72 (Human CDK1) in green with inhibitor bound (in yellow), aligned to *C. elegans* CDK-1 homology model in blue. The human and *C. elegans* kinase domains share 67% identity. All residues that are proximal to the inhibitor binding site with side-chains facing inwards are highlighted on the alignment (identified from PDB structures 4Y72, 6GU4 and associated *C. elegans* homology models). Residues of interest are indicated on the structure above in dark pink (*C. elegans* residue) and light pink (human residue). **(H) Structure and alignment of MAPK14 and PMK-1**. Structure of PDB: 6SFO (Human MAPK14) in green with SR-318 bound (in yellow), aligned to *C. elegans* PMK-1 homology model in blue. The human and *C. elegans* kinase domains share 71% identity. All residues that are proximal to the inhibitor binding site with side-chains facing inwards are highlighted on the alignment (identified from PDB structures 6SFO, 3ZS5, 1DI9, 1KV2 and associated *C. elegans* homology models). Residues of interest are indicated on the structure above in dark pink (*C. elegans* residue) and light pink (human residue). **(I) Structure and alignment of SRC and SRC-1**. Structure of PDB: 6ATE (Human SRC) in green with inhibitor bound (in yellow), aligned to *C. elegans* SRC-1 homology model in blue. The human and *C. elegans* kinase domains share 63% identity. All residues that are proximal to the inhibitor binding site with side-chains facing inwards are highlighted on the alignment (identified from PDB structures 6ATE, 4MXO and associated *C. elegans* homology models). Residues of interest are indicated on the structure above in dark pink (*C. elegans* residue) and light pink (human residue). **(J) Structure and alignment of IGF1R and DAF-2**. Structure of PDB: 5FXS (Human IGF1R) in green with inhibitor bound (in yellow), aligned to *C. elegans* DAF-2 homology model in blue. The human and *C. elegans* kinase domains share 47% identity. All residues that are proximal to the inhibitor binding site with side-chains facing inwards are highlighted on the alignment (identified from PDB structures 5FXS, 4D2R and associated *C. elegans* homology models). Residues of interest are indicated on the structure above in dark pink (*C. elegans* residue) and light pink (human residue). **(K) Structure and alignment of STK10 and GCK-4**. Structure of PDB: 6EIM (Human STK10) in green with GW683134A bound (in yellow), aligned to *C. elegans* GCK-4 homology model in blue. The human and *C. elegans* kinase domains share 56% identity. All residues that are proximal to the inhibitor binding site with side-chains facing inwards are highlighted on the alignment (identified from PDB structures 6EIM, 4EQU and associated *C. elegans* homology models). Residues of interest are indicated on the structure above in dark pink (*C. elegans* residue) and light pink (human residue). **(L) Structure and alignment of CSNK1D and KIN-20**. Structure of PDB: 5MQV (Human CSNK1D) in green with inhibitor bound (in yellow), aligned to *C. elegans* KIN-20 homology model in blue. The human and *C. elegans* kinase domains share 80% identity. All residues that are proximal to the inhibitor binding site with side-chains facing inwards are highlighted on the alignment (identified from PDB structures 5MQV, 5OKT and associated *C. elegans* homology models). Residues of interest are indicated on the structure above in dark pink (*C. elegans* residue) and light pink (human residue). **(M) Structure and alignment of GSK3B and GSK-3**. Structure of PDB: 6HK3 (Human GSK3B) in green with inhibitor bound (in yellow), aligned to *C. elegans* GSK-3 homology model in blue. The human and *C. elegans* kinase domains share 82% identity. All residues that are proximal to the inhibitor binding site with side-chains facing inwards are highlighted on the alignment (identified from PDB structures 6HK3, 5HLN and associated *C. elegans* homology models). Residues of interest are indicated on the structure above in dark pink (*C. elegans* residue) and light pink (human residue). **(N) Structure and alignment of BUB1 and BUB-1**. Structure of PDB: 6F7B (Human BUB1) in green with BAY-1816032 bound (in yellow), aligned to *C. elegans* BUB-1 homology model in blue. The human and *C. elegans* kinase domains share 30% identity. All residues that are proximal to the inhibitor binding site with side-chains facing inwards are highlighted on the alignment (identified from PDB structures 6F7B, 4QPM and associated *C. elegans* homology models). Residues of interest are indicated on the structure above in dark pink (*C. elegans* residue) and light pink (human residue). **(O) Structure and alignment of CSNK2A1 and KIN-3**. Structure of PDB: 3R0T (Human CSNK2A1) in green with inhibitor bound (in yellow), aligned to *C. elegans* KIN-3 homology model in blue. The human and *C. elegans* kinase domains share 86% identity. All residues that are proximal to the inhibitor binding site with side-chains facing inwards are highlighted on the alignment (identified from PDB structures 3R0T, 3NSZ and associated *C. elegans* homology models). Residues of interest are indicated on the structure above in dark pink (*C. elegans* residue) and light pink (human residue). **(P) Structure and alignment of CSNK1A1 and KIN-19**. Structure of PDB: 6GZD (Human CSNK1A1) in green with inhibitor bound (in yellow), aligned to *C. elegans* KIN-19 homology model in blue. The human and *C. elegans* kinase domains share 90% identity. All residues that are proximal to the inhibitor binding site with side-chains facing inwards are highlighted on the alignment (identified from PDB structure 6GZD and associated *C. elegans* homology model). Residues of interest are indicated on the structure above in dark pink (*C. elegans* residue) and light pink (human residue).

**Supplementary Figure 3.**
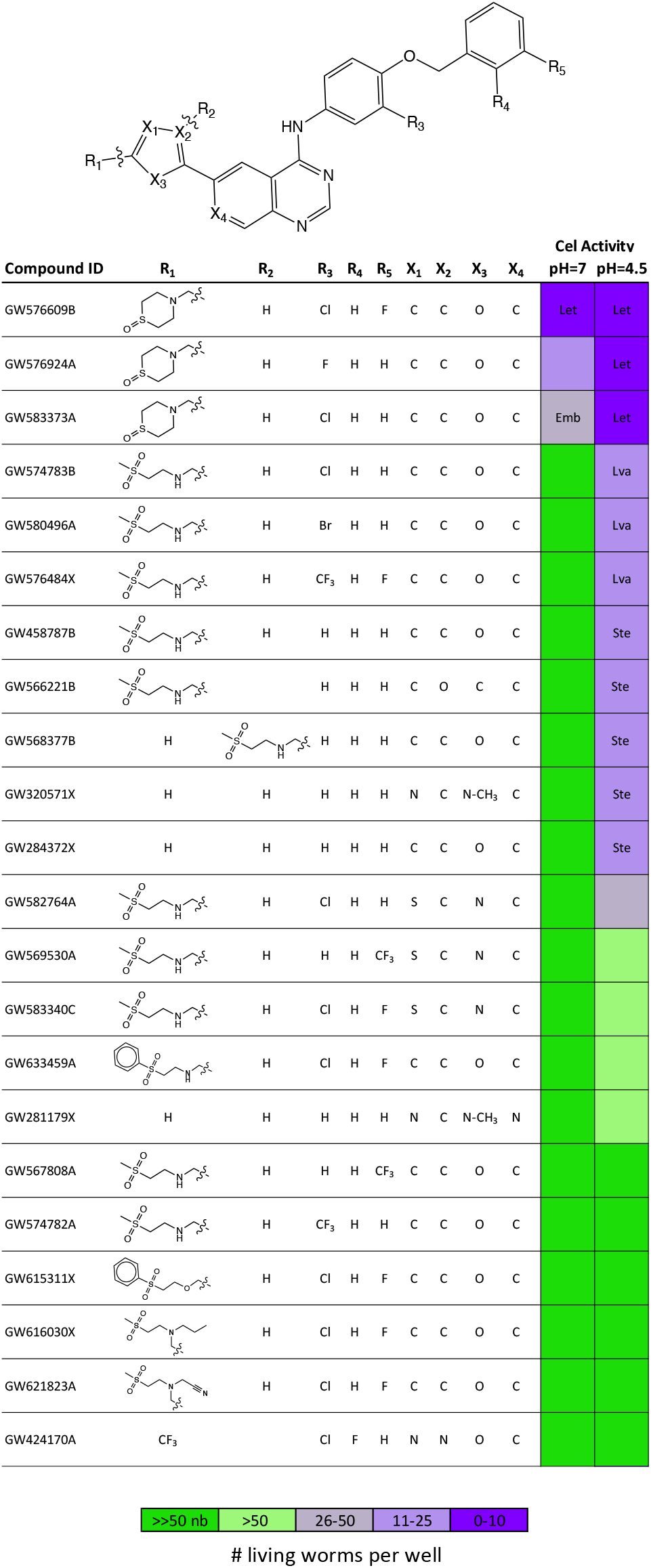
4AQ SAR highlighting favorable structural features for nematode activity *in vivo*. Screen phenotypes induced by inhibitor exposure including lethality (Let), embryonic lethality (Emb), larval arrest (Lva) and sterility (Ste) are shown. The resulting population growth defects are indicated by the colour coded scale (nb, no bacteria remaining in the well).

**Supplementary Figure 4.**
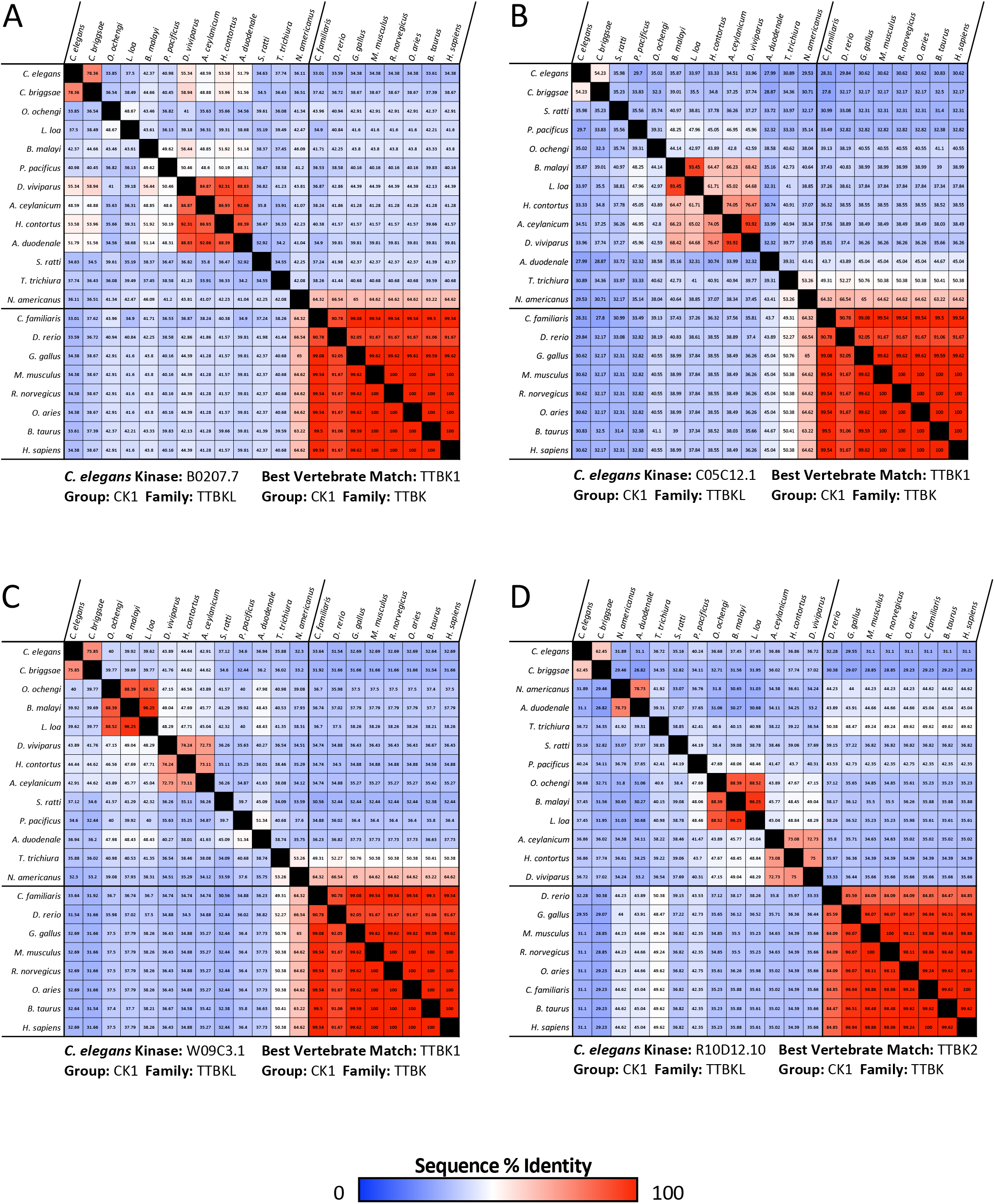
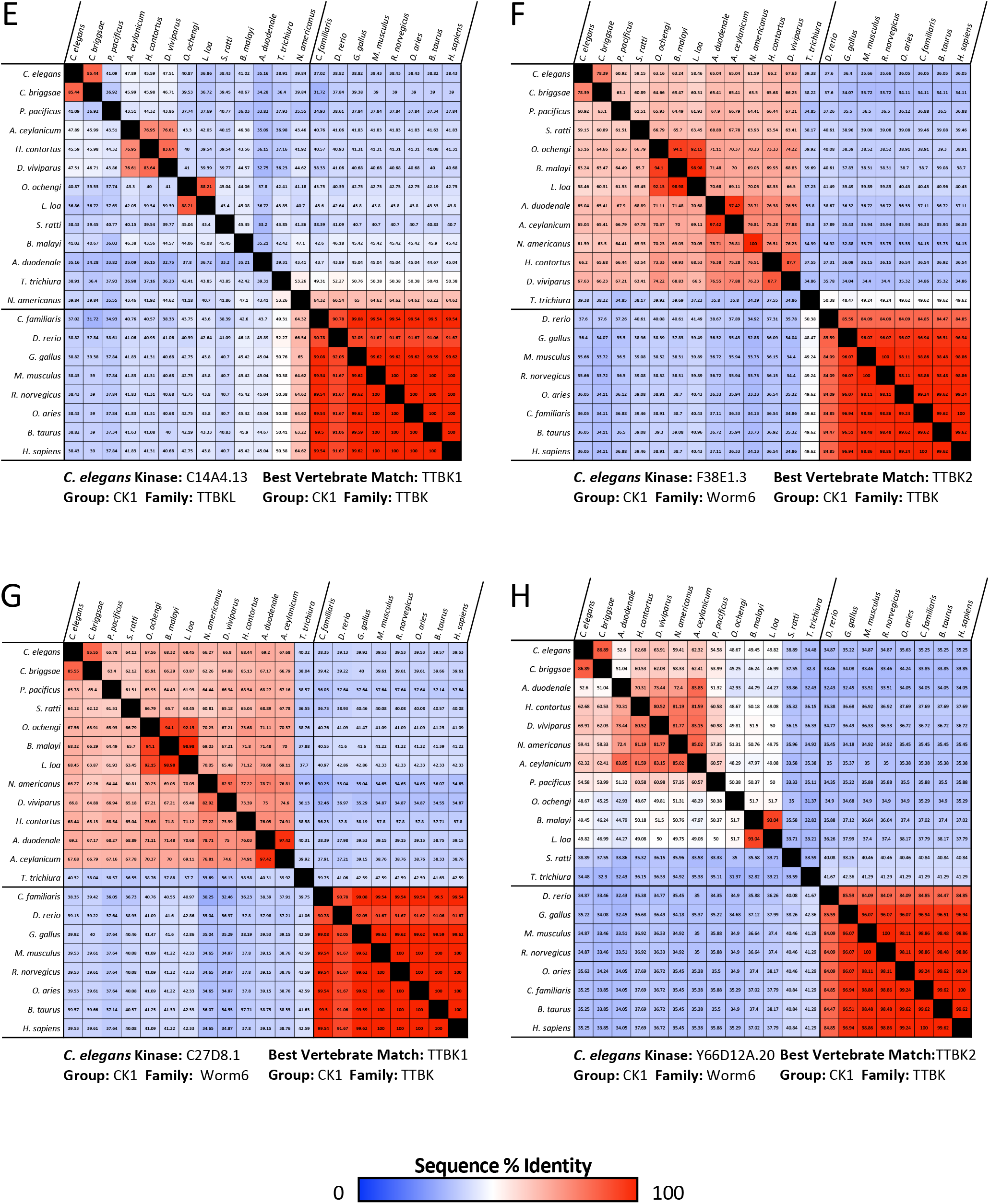
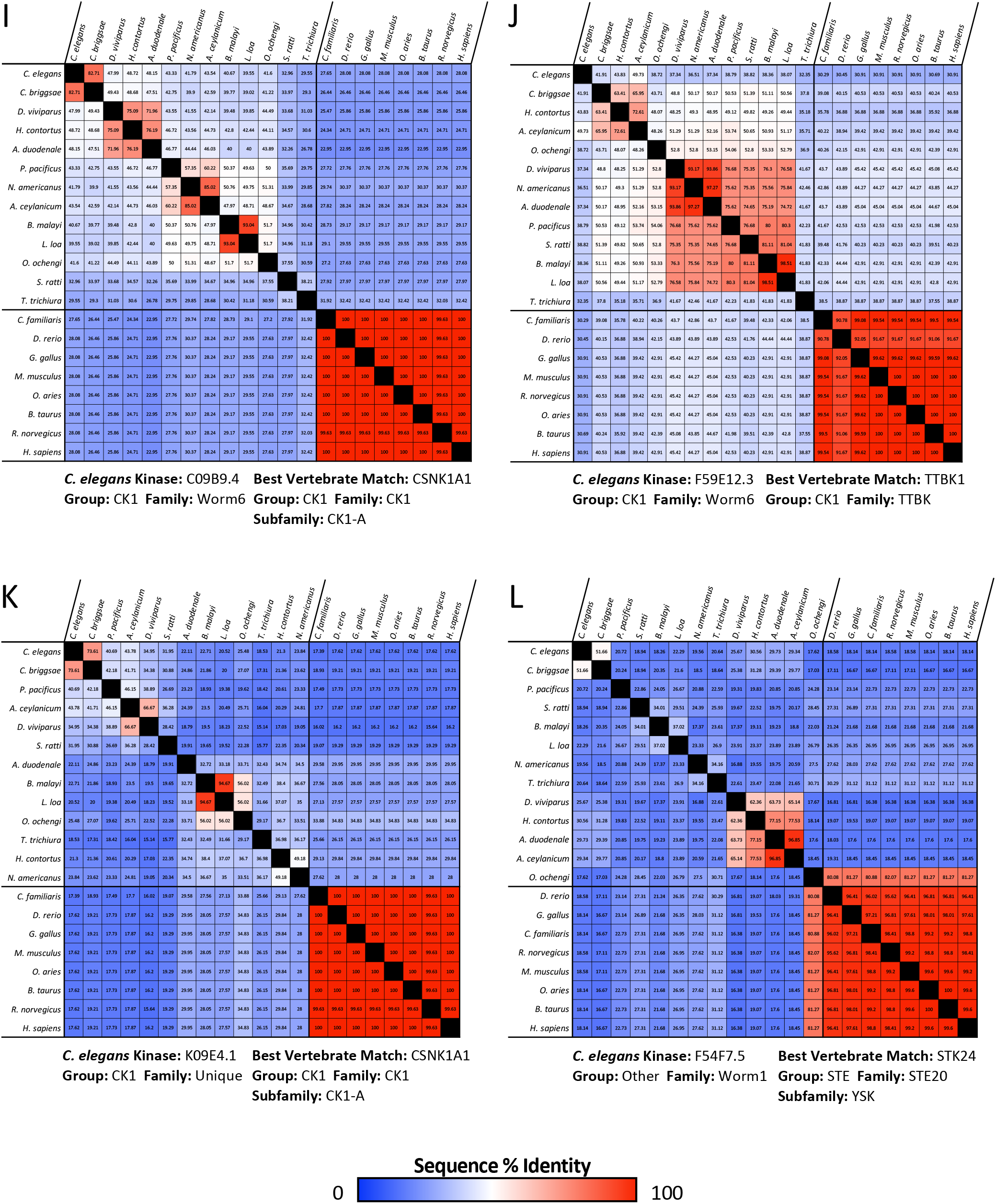
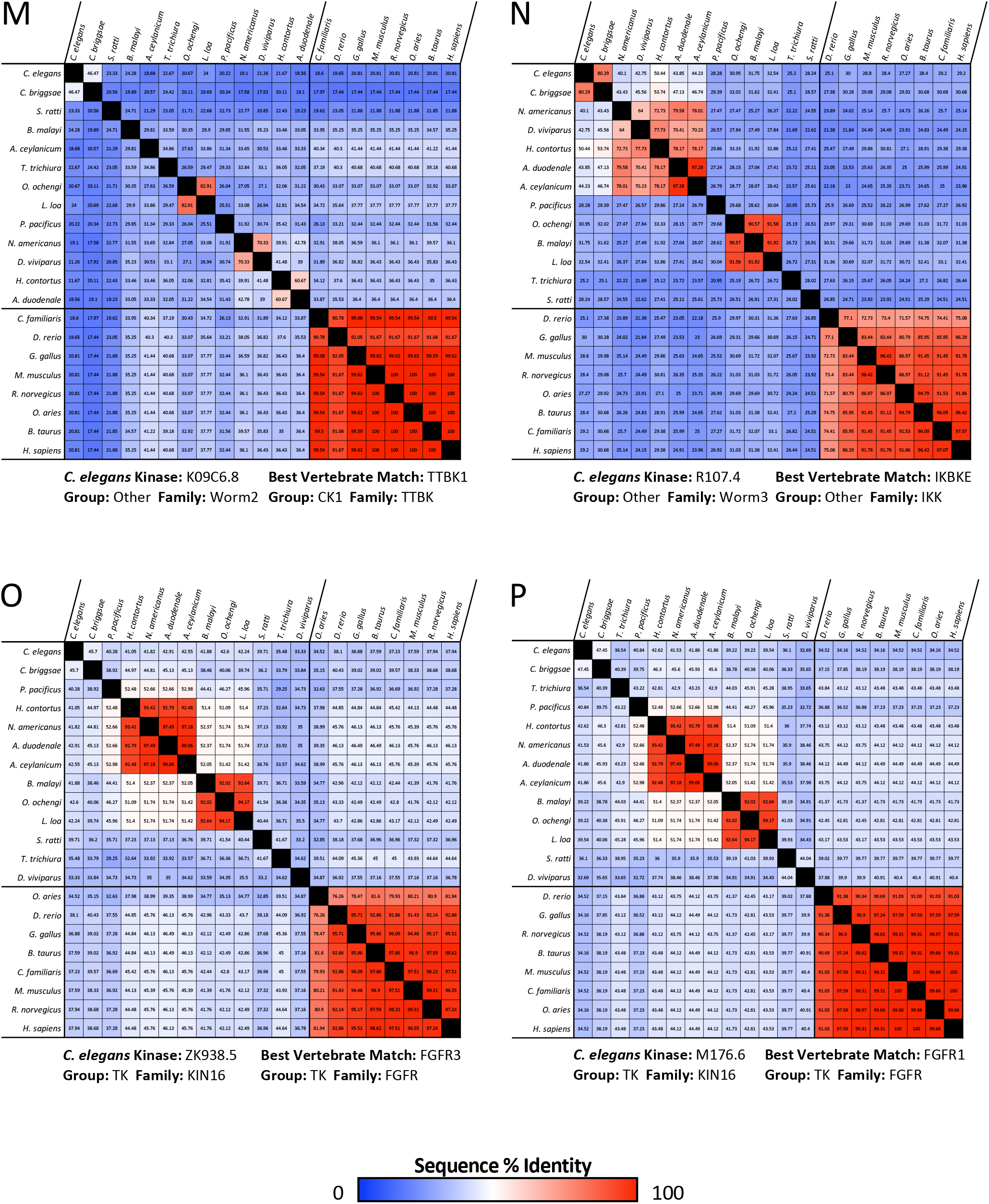
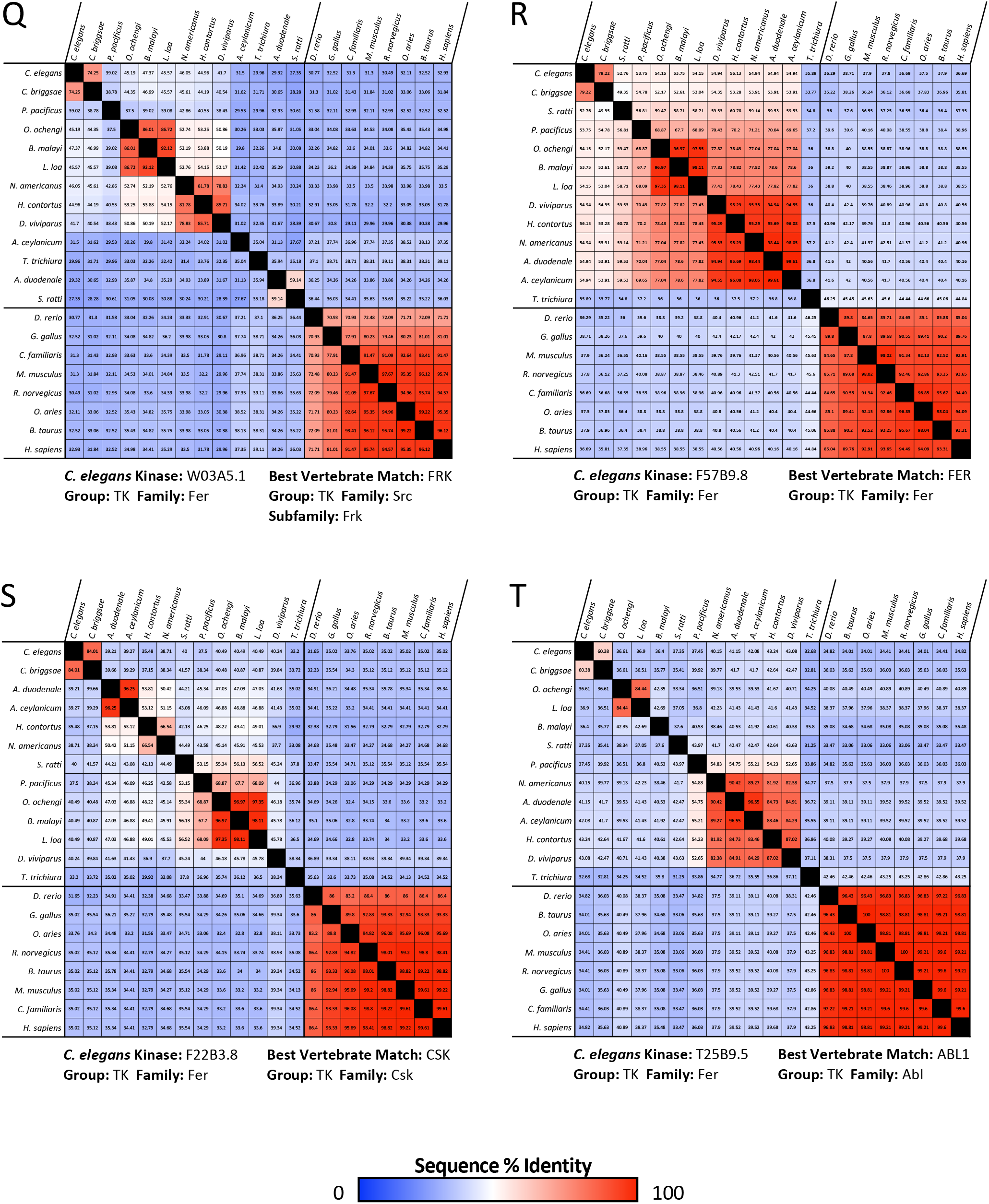
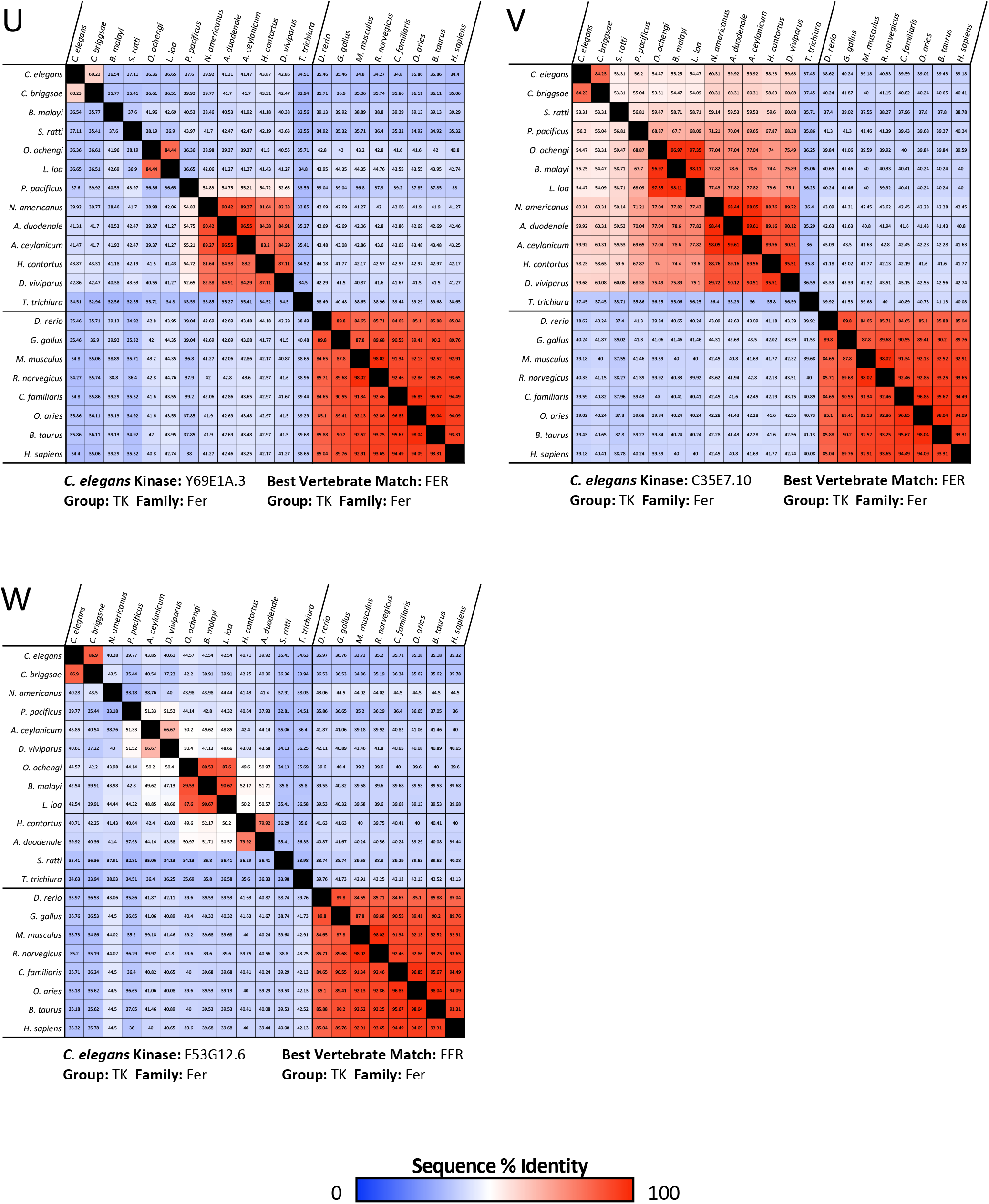
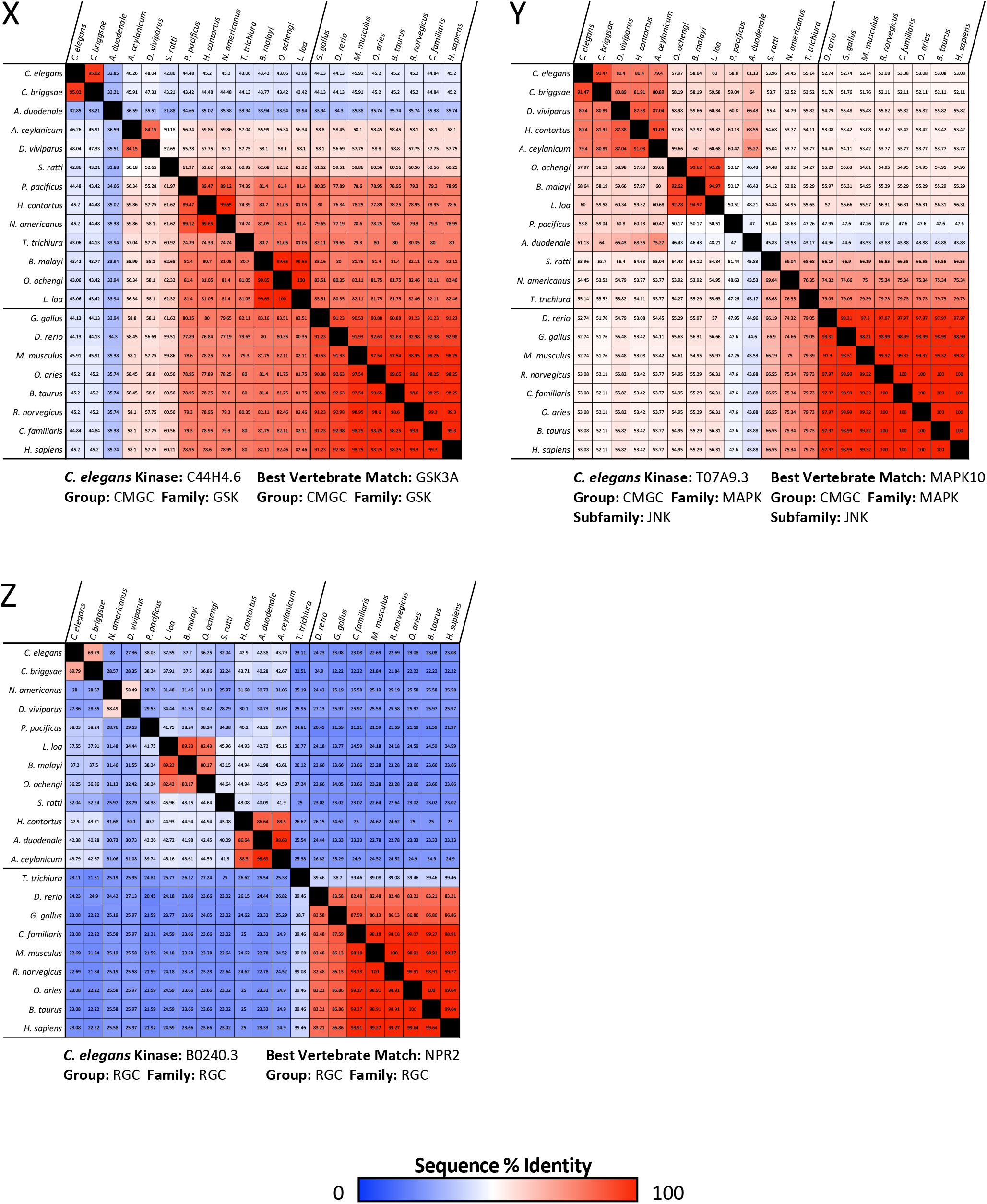
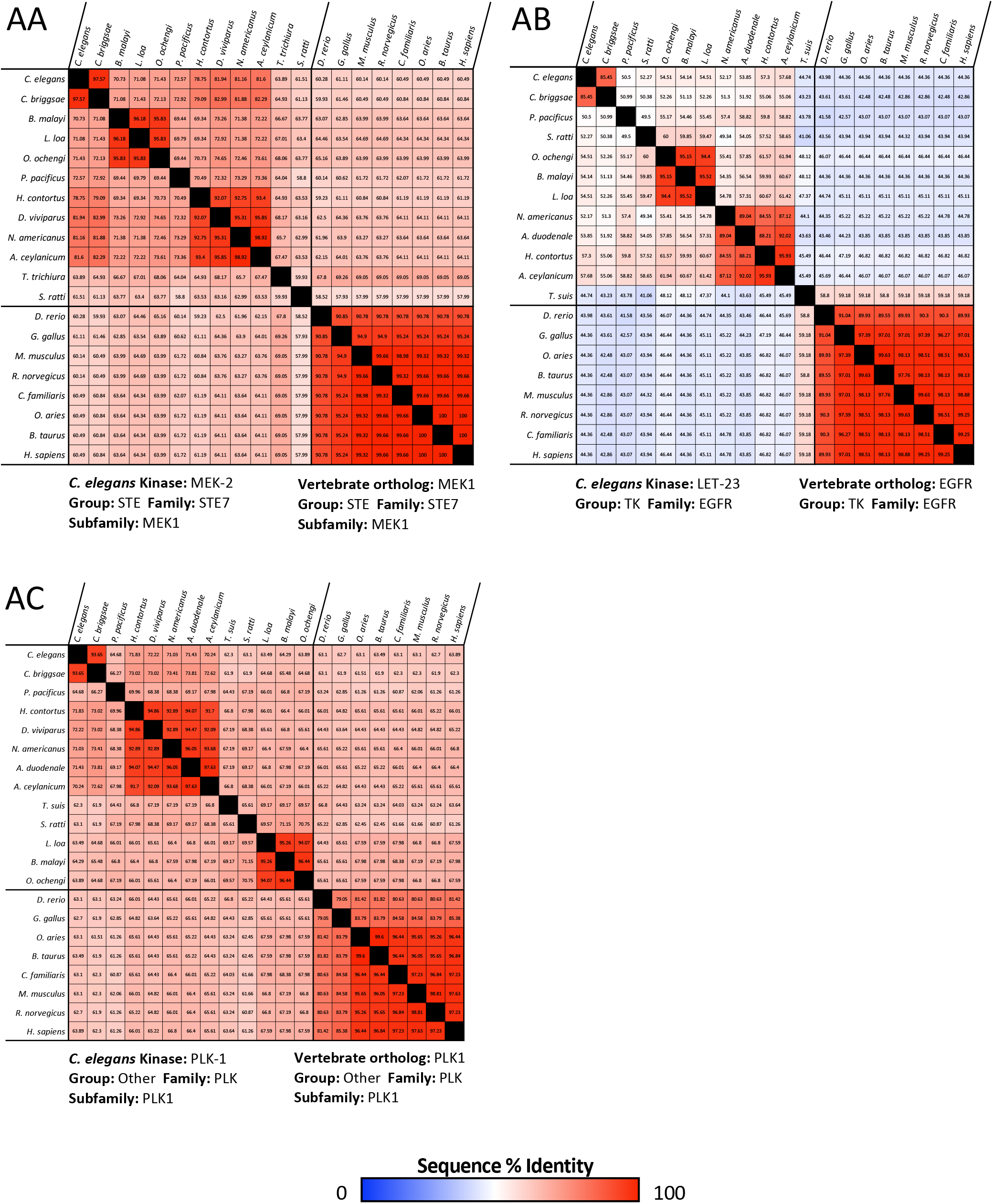
Conservation of essential nematode-specific kinases across species. Matrices show the percent sequence identity between the kinase domain of the *C. elegans* essential nematode-specific kinase and that of the most similar kinase found across nematode species along with the kinase domain sequence of the best human kinase match and its ortholog across vertebrate species. *C. elegans* kinases from nematode-specific families (A-P) and nematode-expanded families (Q-Z) are shown. Sequence identity matrices for the kinase domain of the well-conserved kinases MEK1/MEK-2, PLK1/PLK-1 and EGFR/LET-23 are included for comparison (AA-AC). Kinase sequences were identified using NCBI BLASTP. Percent identity matrices were generated using Clustal Omega.

## Acknowledgements

The Ontario Institute for Cancer Research (OICR) Kinase Inhibitor Library was provided by Rima Al-Awar and David Uehling at OICR. The APExBIO DiscoveryProbe Kinase Inhibitor Library was provided by Jim Dowling. The GSK Published Kinase Inhibitor Set (PKIS) molecules were provided by William Zuercher. We are grateful for the gift of the humanized LET-23 *C. elegans* strain from Dr. Jaegal Shim. Some *C. elegans* strains were provided by the CGC, which is funded by NIH Office of Research Infrastructure Programs (P40 OD010440).

This work is supported by a CIHR grant (313296) and a Canada Research Chair in Chemical Genetics to PJR and an NSERC Alexander Graham Bell Canada Graduate Scholarship to JK.

